# The Locomoting State Selectively Amplifies Activity of Sensitizing Neurons in Primary Visual Cortex

**DOI:** 10.1101/2025.07.24.666602

**Authors:** A.J Hinojosa, Ye Kosiachkin, S.E Dominiak, B.D Evans, L Lagnado

## Abstract

Sensory processing in the cortex reflects both adaptation to external stimuli and changes in internal state. To investigate how these processes interact in layer 2/3 of mouse V1 we combined calcium imaging, optogenetics and circuit modelling. We find that locomotion preferentially increases the responses of pyramidal cells (PCs) that sensitize during visual stimulation compared to those that depress. A model explains this differential modulation through: (i) variations in the strength of PV and SST connectivity to individual PCs, (ii) broad locomotion-dependent weakening of PC and PV synapses, and (iii) reduced SST inhibition targeting sensitizing PCs. Differences in PV:SST input ratios across sensitizing and depressing PCs can be reproduced by a random- connectivity model based on measured interneuron densities, connection probabilities and synaptic strengths. Thus, stochastic variation in local inhibitory connectivity can explain much of the functional heterogeneity across PCs, while state-dependent modulation of inhibitory synapses reorganizes this balance during locomotion to bias cortical population activity towards sensitizing dynamics. The apparently paradoxical combination of increased PC response but decreased synaptic strength is consistent with a state-dependent gating mechanism that boosts signals leaving V1 while simultaneously preventing disruption of the local excitatory-inhibitory balance required for stable computation.

## Introduction

Sensory processing in the cortex is adjusted both by adaptation to the external environment and changes in the internal physiology of the animal associated with behavioural states such as attention, arousal and satiety (Wark et al. 2007, Polack et al. 2013, McGinley et al. 2015, Pakan et al. 2016, Lin et al. 2019). The mechanisms by which external and internal variables interact within the cortical circuitry are not understood. Here we investigate this question by analysing how the change in state associated with locomotor activity impacts adaptation in primary visual cortex (V1) of mice (Adesnik 2017).

The dynamics of neurons in V1 reflect adaptation on multiple time-scales. In pyramidal cells (PCs), the initial excitatory response to an increase in contrast is followed by a fast (subsecond) decrease in gain reflecting depression in feedforward inputs from the LGN (Varela et al. 1997, Chance et al. 1998). This is immediately followed by a slower phase of adaptation that varies dramatically across the PC population: while some PCs depress in response to a high-contrast stimulus others simultaneously *sensitize*, i.e. increase their response amplitude over time (Heintz et al. 2022). PCs within layer 2/3 are not, therefore, a homogenous population: to understand gain control we must take into account variations in local inhibitory circuits in which they are embedded. An important unresolved question is whether this functional heterogeneity requires selective wiring of inhibitory circuits onto different PCs, or whether substantial differences in inhibitory balance can emerge simply from stochastic variation in local connectivity.

The onset of locomotion signals changes in internal state associated with attention or arousal, acting in less than a second to amplify visual responses in PCs (Niell and Stryker 2010, Saleem et al. 2013, Fu et al. 2014, Vinck et al. 2015, Pakan et al. 2016, Dipoppa et al. 2018). A major driver of this increase in gain are cholinergic fibers arriving from the basal forebrain (Goard and Dan 2009, Polack et al. 2013, Pronneke et al. 2015, Lohani et al. 2022). Some of these fibers make direct synaptic connections that excite interneurons expressing vasoactive intestinal peptide (VIP)(Kimura and Baughman 1997), leading to amplification of PC responses through a VIP->SST->PC disinhibitory pathway. The majority of the cholinergic input, however, innervates V1 diffusely to modulate circuit function through extra-synaptic (muscarinic) receptors (Kimura and Baughman 1997, Goard and Dan 2009, Polack et al. 2013). In vitro experiments have shown that M1 and M3 receptors on soma and dendrites of PCs can increase excitability while M2 receptors on axonal compartments reduce synaptic release probability (Hasselmo and Bower 1992, Mrzljak et al. 1993, Kimura and Baughman 1997). Accounting for state-dependent changes in cortical processing therefore requires an integrated understanding of changes in both electrical activity and the strength of synaptic connections (Dipoppa et al. 2018). The difficulty is that modulations of synaptic strengths are very hard to quantify in awake and behaving animals: recording from connected neuron pairs remains technically challenging and is limited to anaesthetized animals in which inhibitory microcircuits are severely compromised (Haider et al. 2013, Jouhanneau et al. 2015, Pala and Petersen 2015). Current understanding of synaptic modulation in cortical circuits is derived from *ex vivo* studies or inferred from indirect *in vivo* observations and modelling (Dipoppa et al. 2018, Keller et al. 2020, Kraynyukova et al. 2022). At the same time, anatomical and electrophysiological measurements show that PV and SST interneurons are relatively sparse and connect to nearby PCs with probabilities substantially below one. These basic features of cortical organization raise the possibility that random connectivity itself generates a broad distribution in the relative PV and SST inhibition received by individual PCs.

To understand how locomotion and adaptation interact in V1 of mice we used calcium imaging to record visual responses in PCs and the three major classes of interneuron (VIP, SST and PV) together with optogenetics to identify key interactions and a data-driven circuit model to estimate changes in synaptic weights. The model demonstrates that long-range inputs to VIPs that increase visual responses in the locomoting state also govern the time-course of adaptation over the behavioural time- scales of seconds. These inputs do not act uniformly across the PC population but preferentially increase the response of neurons that sensitize to a high-contrast stimulus, especially at lower running speeds. Contrary to a simple model of gain control through the VIP->SST->PC pathway we find that visual responses in SSTs are not inhibited by locomotion but instead weakly enhanced. The model reconciles these observations through a neuromodulatory decrease in synaptic outputs across PCs and interneurons. Locomotion preferentially increases the response of PCs that undergo the sensitizing form of adaptation because these receive a stronger SST connection than depressor PCs rendering them more sensitive to the weakening of SST synapses.

We further show that variations in PV:SST input ratio predicted by the model are largely reproduced by simulations in which PV and SST connections are assigned randomly according to experimentally measured cell densities, connection probabilities and synaptic strengths. Random local connectivity can therefore generate much of the inhibitory heterogeneity associated with opposing forms of adaptation, while state-dependent recruitment and changes in synaptic strength reshape this distribution during locomotion. These results provide an integrated view of how local circuit architecture and behavioural state interact to adjust visual processing in V1 when an animal switches between resting and active states.

## Results

### Differential effects of locomotion on adaptation of pyramidal cells

To understand how changes in brain state interact with adaptation in V1 we used two-photon calcium imaging to measure the activity of excitatory and inhibitory neurons in layer 2/3 during periods of rest and locomotion (Fig 1A, Supplementary Fig. 1, see Methods). The GCaMP6f signal in PCs responding to a prolonged, high contrast stimulus (20° drifting grating at 1 Hz, 10 s duration) are shown in Fig. 1B. Locomotion increased the response averaged across all PCs by a factor of 1.9 ± 0.2. The dynamics of these responses varied widely between individual neurons; while some PCs responded with high initial amplitude and then depressed, others responded with low initial amplitude and then sensitized. This heterogeneity is illustrated by the raster plot in Fig. 1B which shows 522 neurons ordered by the time of peak activity: depression (early peak) and sensitization (late peak) were similarly represented in the two behavioural states (Fig 1B, top). Previous work has shown that receptive field location and orientation tuning are not significantly different between depressing and sensitizing PCs (Heintz et al. 2022).

**Figure 1.**
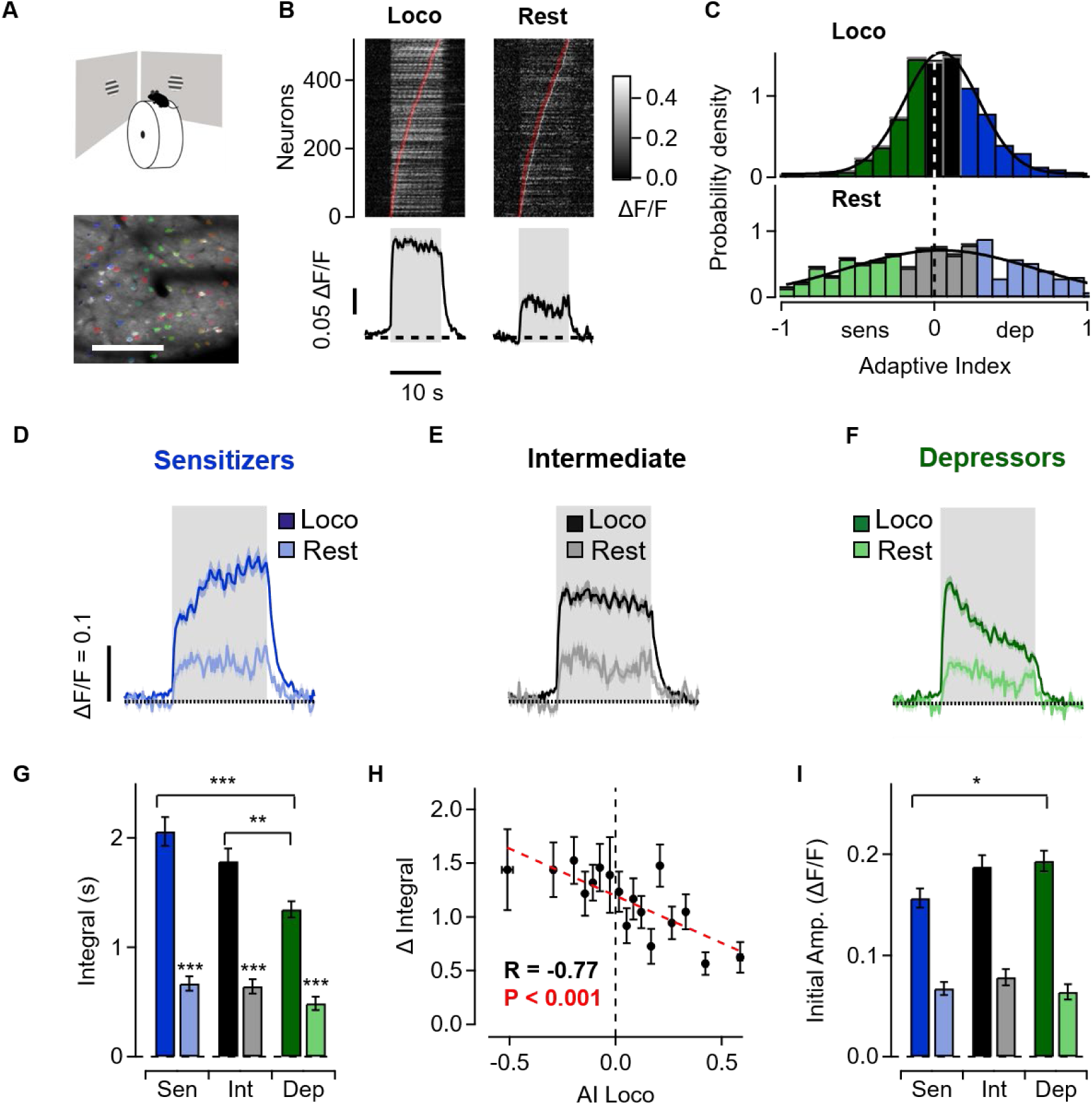
Slow adaptation in pyramidal cells during rest and locomotion. **A.** Top: Mice running on a treadmill were shown a visual stimulus (drifting grating, 20° in size) for 10 s. Bottom: example field of view (FOV) showing PCs in V1 expressing GCaMP6f and corresponding regions-of-interest (ROIs) for individual neurons (coloured). Scale bar 200 µm. **B.** Top: raster plots showing average calcium responses from individual PCs that were responsive to the stimulus during both locomotion and rest (n = 522 neurons, from 9 mice, cells nested within animals). Responses were averaged over 10 stimulus trials then sorted by time of maximum amplitude in rest and locomotion independently. Bottom: average calcium response of all PCs to the visual stimulus during locomotion and rest. **C.** Distributions of adaptive indices (AIs) in the PC population during locomotion (top) and rest (bottom) in sensitizers (blue), intermediate cells (grey) and depressors (green). Both distributions can be described by Gaussians with similar x0 but different width (Rest, x0 =0.04 ± 0.05, width = 0.88 ± 0.08; Loco, x0 = 0.05 ± 0.01, width = 0.34 ± 0.02). Note that AI was calculated separately in this figure only to compare adaptive properties during locomotion and rest. **D, E, F.** Splitting the distribution of PCs according to their AI during locomotion reveals that the tertile with lower AI showed slow sensitization, the tertile with intermediate values showed little adaptation, and the tertile with higher AI showed slow depression. **G.** Differences in response of sensitizers (blue), intermediate cells (grey) and depressors (green) measured as the integral during the stimulus. Locomotion caused a larger increase in sensitizers than depressors (interaction between locomotion state and AI: p < 0.001, Linear Mixed Model). *p < 0.05, **p < 0.01, ***p < 0.001. **H.** Locomotion-induced increase in response integral as a function of AI during locomotion. PCs were ordered by AI and grouped into sets of 30 cells. Correlation was calculated across the mean values of these groups (R = −0.77, p < 0.001). **I.** Initial response amplitude in sensitizing, intermediate and depressing PCs, classified according to their AI during locomotion. Data are mean ± SEM

The increase in response during locomotion was manifested in two ways; a 6-fold increase in the number of PCs generating a significant response (Supplementary Table 1) and an increase in the average response, measured as the integral during the 10 s stimulus (Fig. 1B). To investigate how these response changes affected PCs undergoing different forms of adaptation, we split the population into three equal sized groups based on the distribution of the adaptive index (AI): sensitizing (more negative AI), intermediate, and depressing (more positive AI; Fig. 1C). This was calculated as AI = (R1 − R2)/(R1 + R2) where R1 is the average response over the period 0.5 - 2.5 s after stimulus onset and R2 is the average at 8-10 s (Methods). The AI, when computed separately for rest and locomotion and averaged across the whole PC population, was similar during locomotion (0.05 ± 0.3; mean ± sem) and rest (0.04 ± 0.9; Fig. 1C). The magnitude of the visual response, however, increased by a factor of 2.1 ± 0.4 in sensitizing PCs, 1.7 ± 0.3 in the intermediate group and 1.7 ± 0.4 in depressors. Cells were classified according to their AI during locomotion to preserve cell identity across behavioural states, as AI was only weakly correlated between rest and locomotion (Supplementary Fig. 2A). These differences were due to sensitizers displaying a larger response during the locomotion state, rather than differences in responses during the resting state (Fig. 1 D- I), and could not be explained by differences in initial response amplitude or baseline activity (Fig. 1 I, Supplementary Fig. 2B,D–F). Consistent with this preferential amplification, PCs that became more sensitizing during locomotion also showed larger increases in response integral (Supplementary Fig. 2C). Locomotion therefore produced a significantly greater response increase in sensitizing than in depressing PCs (Fig. 1D- H).

Many studies treat locomotion as a binary behavioural state (Niell and Stryker 2010, Fu et al. 2014, Pakan et al. 2016, Dadarlat and Stryker 2017, Dipoppa et al. 2018), but there is increasing evidence indicating that visual responses are also modulated by running speed (Saleem et al. 2013, Vinck et al. 2015). We found that response amplitude does not simply increase monotonically with speed but rather exhibits a maximum at 10-20 cm/s followed by a second weaker peak at 45-55 cm/s (Fig. 2A). These speed ranges overlap with those typically associated with walking (9 ± 5 cm/s) and trotting (51 ± 9 cm/s) in mice (Bellardita and Kiehn 2015). Sensitizing PCs consistently underwent a larger increase in visual responses than depressors, with the largest difference occurring at speeds of ∼10 cm/s (Fig. 2B,C).

**Figure 2.**
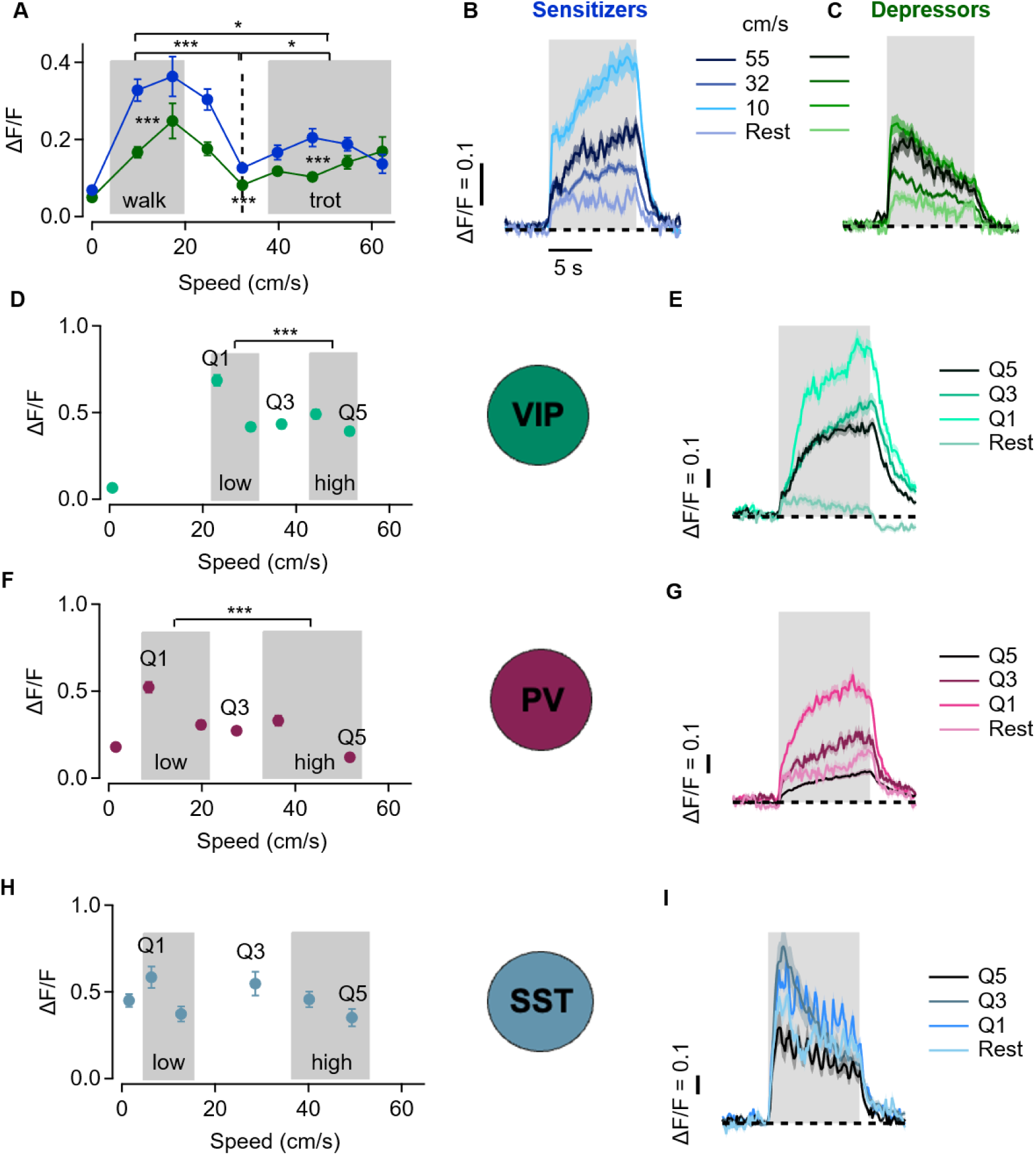
Amplification of visual responses is maximal at fast walking speed. **A.** Locomotion-induced increase in response amplitude (ΔF/F) as a function of running speed in sensitizing and depressing PCs (blue, green). Cells were grouped in speed intervals increasing by 7.5 cm/s (locomotion: 836 cells per PC type from 14 mice; rest: 174 cells per PC type from 9 mice). Shaded regions indicate walking and trotting speed ranges(Bellardita and Kiehn 2015). The two ranges were compared with each other and with the intermediate-speed bin (dashed line); significance is indicated above the graph. Sensitizers and depressors were also compared within each speed range, with significance indicated adjacent to the corresponding range. **B–C.** Average calcium responses of sensitizing (B) and depressing (C) PCs at rest and across different ranges of locomotion speed. **D.** Response amplitude of VIP interneurons as a function of running speed (locomotion: 951 cells; rest: 214 cells; 2 mice). Locomotion data were divided into five equal-sized groups according to running speed (Q, quintile). For statistical analysis, Q1 and Q2 were averaged and compared with the average of Q4 and Q5. **E.** Average calcium responses of VIP interneurons at rest and across three locomotion speed groups shown in D. **F–G.** As in D–E for PV interneurons (locomotion: 669 cells; rest: 171 cells; 3 mice). **H–I.** As in D–E for SST interneurons (locomotion: 327 cells from 6 mice; rest: 115 cells from 3 mice). Horizontal and vertical error bars represent SEM of running speed and response, respectively. Data are shown as mean ± SEM. LMM: * p < 0.05, ** p < 0.01, *** p < 0.001.

Locomotion has also been shown to modulate inhibitory interneurons(Dipoppa et al. 2018), but the impact on their visually-evoked responses is less well characterised. We found that visual responses of VIP and PV interneurons were strongly amplified at lower locomotion speeds (Fig. 2D-G), while SSTs were not significantly affected (Fig.2 H,I). Overall, these results reveal a preferred speed at which locomotion most strongly modulates visual processing in layer 2/3, with the strongest effects in sensitizing PCs.

### Differential effects of locomotion on fast and slow components of adaptation in PCs and interneurons

The immediate spiking response to a high-contrast stimulus is dominated by depressing adaptation (Varela et al. 1997, Chance et al. 1998) but the raw signal from calcium reporter proteins such as GCaMP does not easily distinguish this phase because of the gradual accumulation of calcium in the neuron. Several algorithms for estimating spike rates from calcium signals are now in use (Deneux et al. 2016, Friedrich et al. 2017, Rupprecht et al. 2021) and we applied MLspike because it is based on a biological model of GCaMP activation and calcium dynamics that has been shown to perform better than others (Berens et al. 2018) (Fig. 3A-C). Despite the limited time-resolution of calcium imaging, the inferred spiking response revealed the initial phase of fast depressing adaptation that is a consistent feature of electrophysiological recording (Varela et al. 1997, Chance et al. 1998, Fairhall et al. 2001) (red arrows In Fig. 3A-B). The initial peak was also recovered using the CASCADE (Rupprecht et al. 2021) spike inference method (Supplementary Fig. 3C).

**Figure 3.**
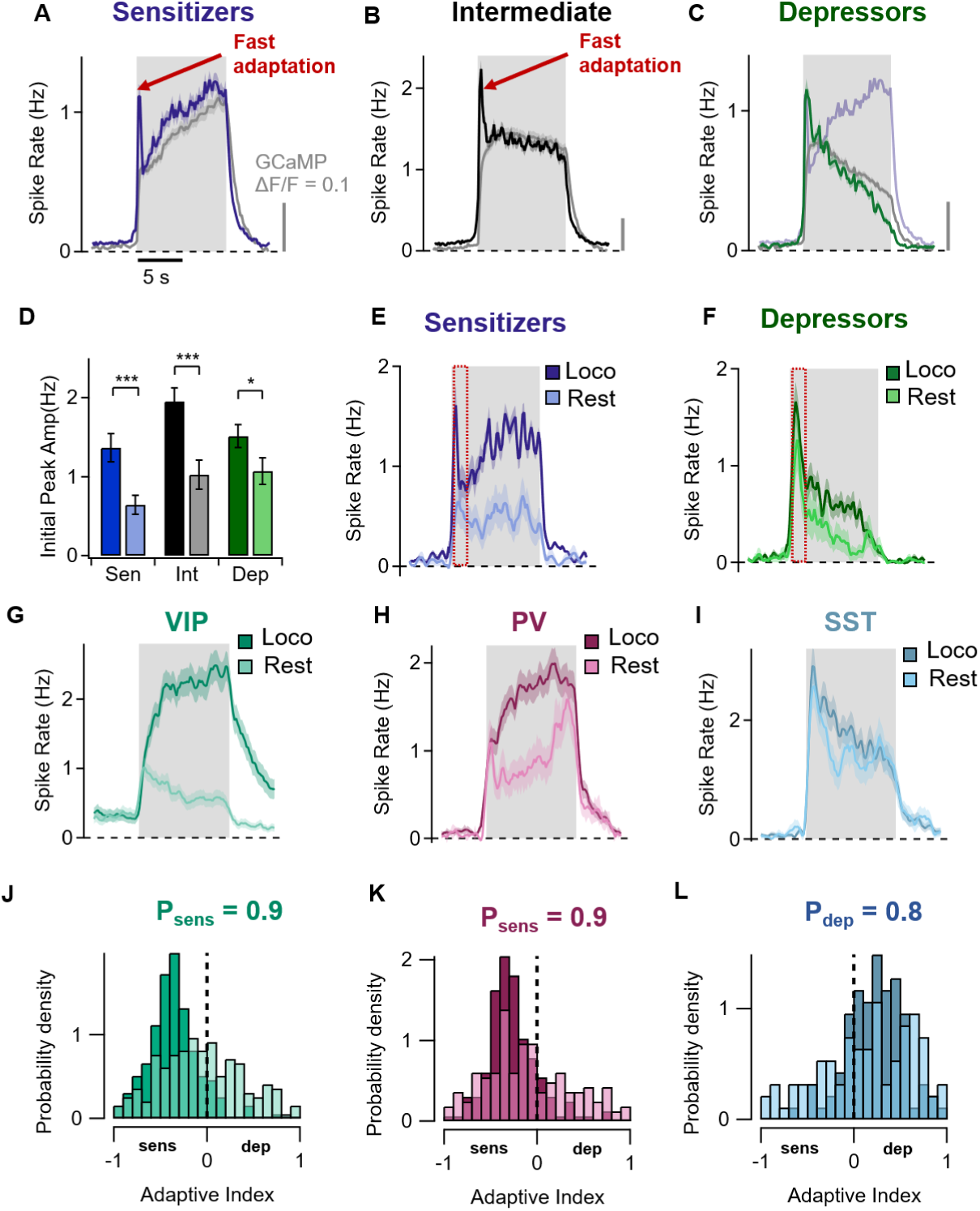
Differential effects of locomotion on fast and slow components of adaptation in PCs and interneurons. **A.** Estimated spike rate averaged across the sensitizing tertile measured during locomotion (Data from all neurons responsive to the stimulus during locomotion, including non-responsive cells during rest; n = 479 neurons from 14 mice). The GCaMP signal averaged across the same set of neurons is shown in grey. The fast depressing component of adaptation is highlighted by the red arrow. **B.** As A, but for the second tertile of PCs that were pre-dominantly non-adapting. **C.** As A, but for the depressing tertile. The trace from sensitizers in A is superimposed (light blue) showing the similar initial response in sensitizing and depressing PCs. **D**. Initial peak amplitude at rest in sensitizing, intermediate and depressing PCs, classified according to their AI during locomotion. **E.** Comparison of average spiking response at rest and during locomotion in the sensitizing tertile of PCs in which paired measurements were made (158 neurons in 9 mice). The relative change in the response during locomotion was 3.1 ± 0.8. **F.** Comparison of average spiking response at rest and during locomotion in the depressing tertile of PCs. The relative change in the response during locomotion was 1.4 ± 0.2 which was significantly smaller than the sensitizing tertile (LMM, p < 0.001; Supplementary Fig. 3B). **G-I.** Average inferred spike rate of VIPs, PVs and SSTs during locomotion and rest. The integrated response during locomotion increased by factor of 5.1 ± 1.0 in VIPs and 1.7 ± 0.2 in PVs but was not significant in SSTs (1.2 ± 0.2). **J-L.** Distribution of adaptive indices in VIPs, PVs and SSTs. AI was measured from GCaMP signals, as for PCs in Fig. 1. VIPs and PVs were predominantly sensitizing (VIPs, Psens = 0.56 at rest and 0.90 during locomotion, n = 122 neurons in 2 mice; PVs, Psens = 0.64 at rest and 0.86 during locomotion, n = 97 neurons in 3 mice). SSTs were predominantly depressing at rest and during locomotion (Psens = 0.37 at rest and 0.21 during locomotion, n = 82 neurons in 4 mice). Results from D to L are from cell-paired recordings in the two states.

During locomotion, the inferred initial peak amplitude was not significantly different in PCs that depressed (1.52 ± 0.14 Hz) or sensitized (1.37 ± 0.18 Hz), consistent with similar strength of the feedforward input across the PC population (Fig. 3C,D). We therefore used inferred spikes to better represent response dynamics, while retaining calcium signals for the population-level analyses in Figs. 1 and 2 because the discrete nature of inferred spikes produces less continuous estimates of adaptive properties, particularly in sparsely firing cells (Supplementary Fig. 3A). The main effects of locomotion on adaptive properties and their dependence on running speed were preserved in the inferred spikes (Supplementary Fig. 3B,D,E).

Comparing activity at rest and while running revealed a powerful effect on both first and second phases of the response but this varied across the PC population. In the sensitizing tertile the initial peak underwent a relative increase of 1.1 ± 0.5 and the response integrated over the whole stimulus period a relative increase of 3.1 ± 0.8 (Fig. 3D,E; Supplementary Fig. 3B). But in the depressing tertile, locomotion only changed the initial fast response by a factor of 0.4 ± 0.3 while the integrated response only increased by a factor of 1.4 ± 0.2 (Fig. 3D,F; Supplementary Fig. 3B). These values differed quantitatively from those measured from calcium signals (Fig. 1), particularly in depressing PCs, but both measures showed a substantially larger locomotion-associated response increase in sensitizing than depressing PCs. Changes in internal state associated with locomotion do not, therefore, act uniformly across the PC population but preferentially increase the responses of neurons that undergo the sensitizing form of adaptation.

### A data-driven model of contrast adaptation and the effects of locomotion

To understand how an active behavioural state differentially alters neural responses across PCs we constructed a population model of signal flow based on neuronal input- output functions expressed as firing rates (Wilson and Cowan 1972, Miller and Troyer 2002, Rubin et al. 2015) (see Methods). The model took into account three aspects of circuits in layer 2/3: *i)* the measured response dynamics of PCs and the three major types of interneurons (Fig. 3G-L); *ii)* the dynamics of signals that arrive from other brain regions (Fig. 4A), and *iii)* the known connectivity patterns of neurons within this layer (Fig. 4B). Code and example data sets are available at https://github.com/lagnadoLab/CortexModel.

**Figure 4.**
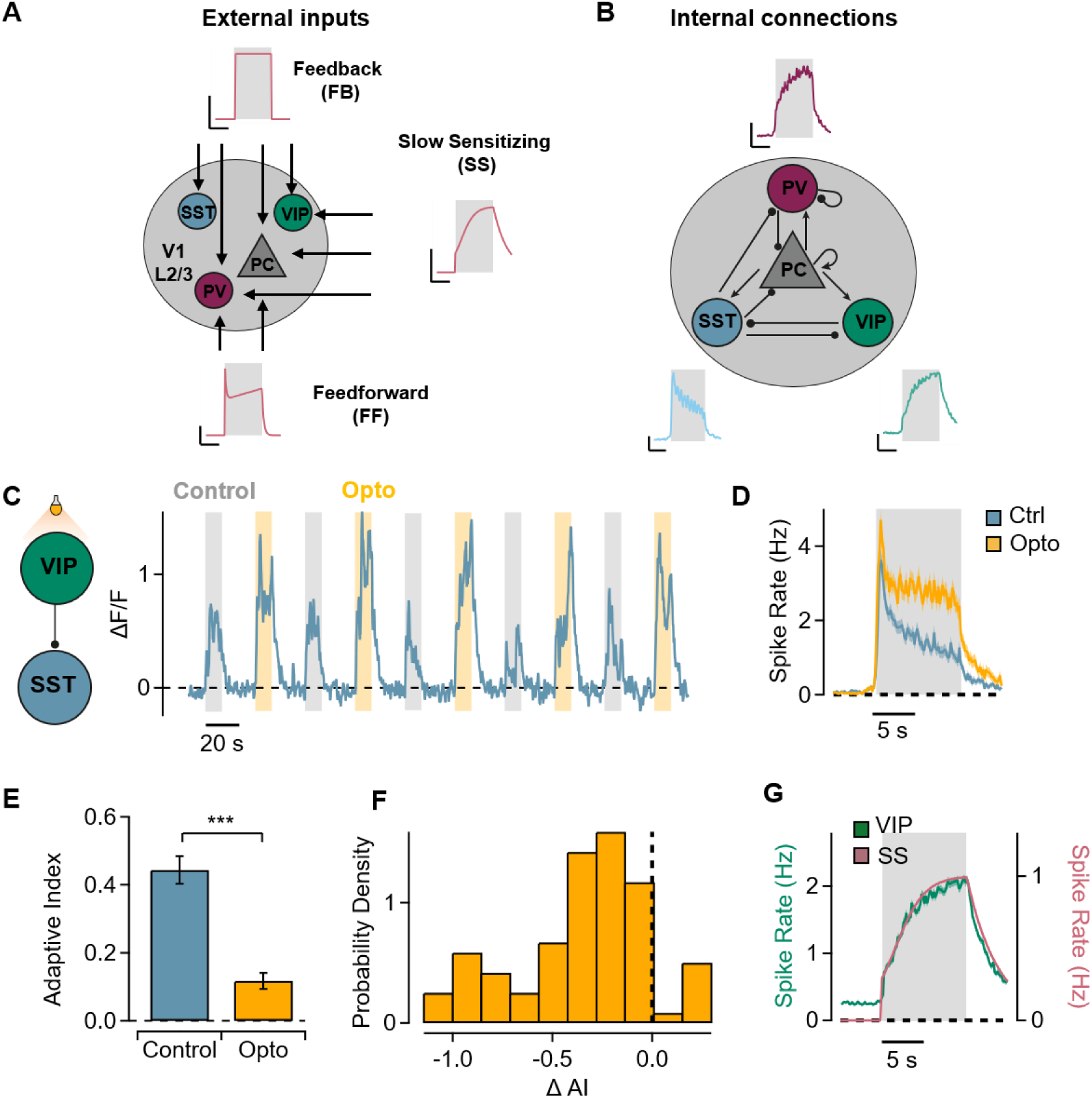
A model of signal flow in layer 2/3. **A.** Schematic showing the targets of three external inputs into layer 2/3 of V1. Feedforward excitation targets PVs and PCs; feedback input targets all populations(Shen et al. 2022, Liu et al. 2024) and the slower sensitizing input enters through VIPs as well as directly activating PCs and PVs(Jordan and Keller 2020) (vertical bar, 0.5 Hz; horizontal bar, 5 s). **B.** Schematic showing the main excitatory (arrow tip) and inhibitory (round tip) connections between cell types within L2/3 of V1 considered in the model. PCs form excitatory connections with all other neuron types including themselves. PVs mainly target PCs and other PVs while SSTs inhibit all other cell types but avoid inhibiting each other. VIPs almost exclusively inhibit SST interneurons (vertical bar, 0.05 ΔF/F; horizontal bar, 5 s). **C.** Left: schematic of the experimental paradigm testing whether VIP inhibition contributes to slow depression in SSTs. VIPs were silenced optogenetically while simultaneously recording activity in SSTs. Right: example response of an SST neuron during stimulus presentation alone and stimulus paired with optogenetic silencing of VIPs during locomotion. **D.** Average response of SSTs with and without optogenetic silencing of VIPs during locomotion (n = 83, from 4 mice). **E.** Adaptive index (AI) of SSTs with and without optogenetic silencing of VIPs. VIP silencing significantly reduced SST AI (linear mixed model, *p* < 0.001). **F.** Distribution of the within-cell change in AI following VIP silencing. AI was reduced in 90% of SSTs. **G.** The slow sensitizing input to the model (SS) was based on the average response of the VIP population during locomotion and consists of a step followed by a sigmoid function with a time-constant = 1.71 s.

#### i) The dynamics of neurons in layer 2/3

A key determinant of the magnitude and direction of slow adaptive changes in a PC is the local inhibitory microcircuit in which it is embedded: PCs in which direct SST inputs dominate over PV inputs tend to sensitize while those in which PV inputs dominate depress (Heintz et al. 2022). We therefore also experimentally measured the effects of locomotion on responses and adaptation in the three major interneuron types to use their response dynamics in the model (Fig. 3G-L; Supplementary Fig. 2G-L). Unlike PCs, interneurons were not divided into tertiles but were classified as sensitizing or depressing according to whether their AI was negative or positive, respectively. VIPs underwent the strongest modulation: the density of responsive neurons changed by a factor of 3 (Supplementary Table 1), and the integrated amplitude of these responses by a factor of 5.1 ± 1.0 (Fig. 3G). Simultaneously, locomotion caused the ratio of sensitizing to depressing VIPs to increase from 1.3 at rest to 9 (Fig. 3J). PVs were more weakly modulated, with integrated responses changing by a factor of 1.7 ± 0.2 while the ratio of sensitizers to depressors also increased from 1.8 to 6.1 (Fig. 3H and K). In contrast, locomotion did not significantly alter the number of responsive SST neurons or the amplitude of their responses (response increase factor of 1.2 ± 0.2; p = 0.28; Fig. 3I and L). This observation was surprising given that VIPs powerfully inhibit SSTs and locomotion activates a VIP->SST->PC disinhibitory pathway often highlighted as an important contributor to gain control(Pfeffer et al. 2013, Fu et al. 2014, Karnani et al. 2016). However, the influence of this pathway is known to vary, consistent with reports that SST response to locomotion depends strongly on the visual stimulus (Pakan et al. 2016, Dipoppa et al. 2018). A major aim of the model, therefore, was to take a more integrated view of gain control that considered multiple interacting pathways within layer 2/3.

The distribution of response dynamics in interneurons was more homogeneous than in PCs. PCs displayed heterogeneous adaptive dynamics spanning sensitization to depression (Fig. 1C), but during locomotion sensitization dominated in VIPs and PVs (P_sens_ = 0.9) while depression dominated in SSTs (P_dep_ = 0.8; Fig. 2J-L). These more consistent, cell-type-specific dynamics allowed us to represent each interneuron class by its characteristic average response (Fig. 3G-I), which we used as empirical input to a population mean-field model. Mean-field models have been widely used to describe the average behaviour of large populations of interacting neurons without simulating each neuron individually (Wilson and Cowan 1972, Brunel 2000, Deco et al. 2008). The network was represented by a system of four first order ordinary differential equations(Wilson and Cowan 1972):

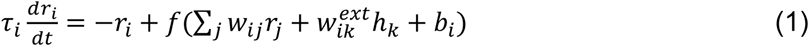

where *i*, *jj* are iterating through the four neuronal populations (PC, VIP, PV and SST), *jj* is a presynaptic population and *i* postsynaptic, *r* is a firing rate and τ_i_ is the time-constant for that population. *f*(*x*) is a neuronal input-output function and here we used a rectified power law (Rubin et al. 2015). The terms 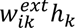 represent the external inputs, where ℎ*_k_* is the activity of external input *k* and 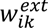 its connection weight onto postsynaptic population *i*. The term *b_i_* is the baseline activity of postsynaptic cell *i*. By fitting the model, we estimated the *connection weight* from presynaptic population *j* to postsynaptic population *i, w_iii_*. This is the product of the total number of synaptic connections from *j* to *i*, *C_iii_*, and the average strength of an individual synapse, *s_iii_*, which we term *synaptic weight*:

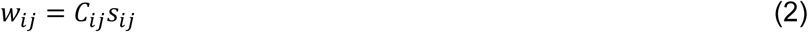

When calculating *s_iii_*, it was important to consider that not all presynaptic neurons are activated by the stimulus and, even if they are, they do not all activate the postsynaptic population because the connection probability (*p_iii_*) is usually less than one. For instance, the average probability of a nearby PV interneuron forming a connection onto a PC is only ∼0.37 at distances less than ∼150 μm, while the probability of an SST interneuron connecting onto a nearby PC is lower, at ∼0.27 (Jiang et al. 2015, Campagnola et al. 2022, Hage et al. 2022). The expected number of connections is therefore *C_iii_* = *N_ii_p_iii_*, where *N_ii_* is the number of responsive presynaptic cells and *p_iii_* the connection probability from cell *j* to cell *i*. Substituting this into equation 2 gives:

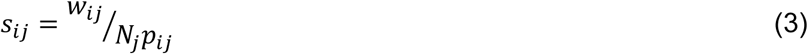

Below we infer changes in individual synaptic weights (*s_iii_*) associated with a transition into the active state by comparing experimentally measured changes in *N_ii_* (Supplementary Table 1) to changes in connection weights (*w_iii_*) estimated by fitting the model.

#### ii) The dynamics of signals entering layer 2/3

Excitatory signals entering layer 2/3 are of three main types (Oh et al. 2014, Froudarakis et al. 2019): feedforward (FF) drive originating from the lateral geniculate nucleus (LGN) and transmitted via layer 4, feedback from higher order cortices (FB) (Roelfsema et al. 1998, Leinweber et al. 2017) and modulated inputs (SS), such as those gated by cholinergic signals from the basal nuclei (Fu et al. 2014). Electrophysiology has shown that FF inputs to layer 2/3 are dominated by fast depressing adaptation occurring within a few hundred milliseconds and the fast phase of adaptation in PCs fell within this range (Varela et al. 1997, Chance et al. 1998) (Fig. 3A-B). Calcium imaging *in vivo* has shown that drifting gratings lasting seconds can also generate a slower sensitizing component in FF inputs that enter layer 2/3 from layer 4 (Makino and Komiyama 2015, Keller et al. 2020). The FF input was therefore modelled with both these components, as shown in Fig. 4A.

The second type of input to V1 is feedback from higher visual areas (Zhang et al. 2014, Fiser et al. 2016, Jin and Glickfeld 2020) and SST interneurons are one of their targets (Adesnik et al. 2012, Zhang et al. 2014, Keller and Mrsic-Flogel 2018, Shen et al. 2022). We have limited understanding of the dynamics of the FB signal over the time- scale of our stimulus, so, for simplicity, it was modelled as a step beginning at a delay of 370 ms after stimulus onset (Fig. 4A), corresponding to the onset of the delayed SST response in our data (Fig. 3I). This is consistent with FB signals emerging after the initial FF response (Kirchberger et al. 2023).

The third type of inputs was required to account for the slow adaptive dynamics observed across the network. This could reflect depressing SST activity or sensitizing activity in VIP and PVs. Because VIPs strongly inhibit SSTs (Pfeffer et al. 2013, Fu et al. 2014, Karnani et al. 2016), we tested whether the depressing dynamics of SSTs depended on VIP activity. We suppressed VIPs while recording SST responses, by expressing the inhibitory photoprotein ArchT in VIPs and GCaMP6f in SSTs (Fig. 4C). Expressing different proteins in two interneuron populations was achieved by crossing a recombinant mouse line expressing Cre (VIP-Cre) with a Flippase expressing line (SST-Flp), then injecting two viral vectors that were Cre and Flippase dependent respectively (see Methods). Suppressing activity in VIPs almost completely abolished the slow component of adaptation in SSTs (Fig. 4D-F). Thus, the depressing dynamics of SSTs depended on activity in the upstream VIP pathway rather than providing an independent source of slow adaptation.

The striking transition from weak, relatively flat VIP responses at rest to strong sensitizing responses during locomotion (Fig. 3G) motivated the inclusion of a sensitizing input to VIPs. We therefore defined the dynamics of this signal from the visual response of VIPs measured experimentally during locomotion (Fig. 4G). We did not include a direct SS input to SSTs because their slow adaptation could be explained by the SS input acting indirectly through VIPs. Visually-driven inputs to VIPs include local recurrent excitation (Pfeffer et al. 2013) and inputs from higher visual areas (Zhang et al. 2014), but the anatomical origin of the SS signal is not established. Its temporal dynamics were therefore modelled empirically from the VIP response rather than assigned to a specific anatomical source (Fig. 4G). This slow sensitizing signal appears quite distinct from inputs from the basal forebrain which provide a binary locomotion state signal but have been found *not* to respond to visual stimuli (Yogesh and Keller 2024). Notably, the dynamics of sensitization in VIPs were similar to sensitization in PCs and PVs suggesting that this input dominated slow adaptation throughout the network (Supplementary Figure 3F,G).

#### iii) Connectivity within layer 2/3

Cortical connections have been probed using a number of approaches, including viral tracing (Kim et al. 2016), electrophysiology (Pfeffer et al. 2013, Jiang et al. 2015, Campagnola et al. 2022) and optogenetics (Hage et al. 2022). Where connectivity differed across studies, we simplified the network by including connections that have been consistently observed in layer 2/3 and are strong enough to significantly influence the circuit (Fig. 4B). For example, PV→SST connectivity has been reported inconsistently across studies and was therefore not included(Pfeffer et al. 2013, Jiang et al. 2015). SST interneurons do not interact with each other while the very small number of connections from VIPs to PCs were neglected because these are not strong enough to significantly influence the circuit (Pfeffer et al. 2013, Hage et al. 2022).

### Using dimensionality reduction and optogenetic manipulations to constrain model solutions and test its predictive ability

The model was fit to average populational responses of PCs and interneurons by adjusting the connection weights *w_iii_* of external and internal connections in layer 2/3 (see Methods). Given the large parameter space spanned by these weights in the model, many solutions were of similar quality. Rather than identifying a single optimal solution, our fitting procedure revealed regions of parameter space that yielded good fits. First, we sampled the parameter space globally by generating 100,000 sets of evenly distributed initial parameters using Sobol sampling (Tennoe et al. 2018, Billeh et al. 2020, Keller et al. 2020). Of these, only a small fraction (∼ 0.03%) produced solutions with reduced chi-square 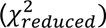 low enough to yield a close fit (rest: 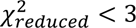; locomotion: 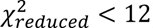), identifying candidate regions of parameter space capable of reproducing the observed responses. We then used these sets to initialise regional sampling, generating bigger populations of fitted solutions for rest and locomotion. To simplify the model, we then examined which parameters differed consistently between behavioural states using effect sizes and principal component analysis. This allowed 10 parameters to be constrained to the same values at rest and during locomotion without substantially reducing the quality of either fit (Supplementary Table 2; See methods).

We then applied a third stage involving a series of experiments in which interneuron activity was altered optogenetically, which further constrained the viable region of parameter space for the locomotion condition (Fig. 5). This third stage could not be usefully applied to data at rest because there were not enough recording episodes across all four optogenetic conditions to provide reliable traces for fitting after subdivision into the four optogenetic conditions (Horrocks et al. 2024). Consequently, weights inferred at rest were less strongly constrained than those inferred during locomotion. In the locomotion condition, the distribution of parameter values after reducing the number of good solutions through all the steps are used to analyse the predictions of the model in Figs. 6 and 7.

**Fig. 5.**
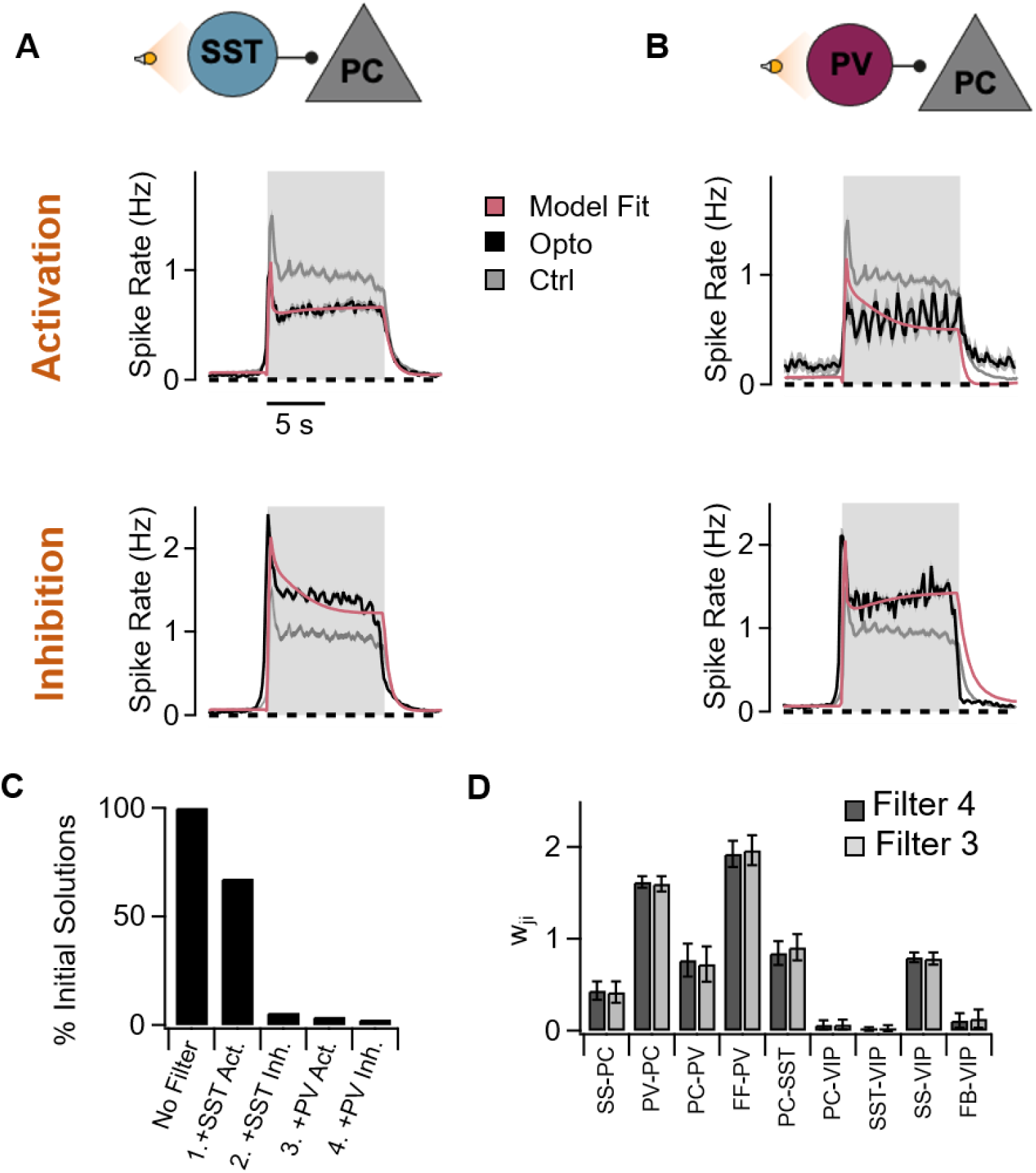
Testing the model using optogenetics. **A.** Top: Average PC response with and without optogenetic activation of SSTs (black and grey). Separate experiments indicated that the intensity of photoactivating light increased SST activity by a factor of ∼1.7(Heintz et al. 2022). The red trace shows the model prediction when SST activity within the model was increased by the same factor. Activation of SSTs decreased PC response and increased sensitization (top, RMSE = 0.05). The grey control trace is averaged over all four series of experiments testing different optogenetic manipulations in A and B. Bottom: Average PC response with and without optogenetic inhibition of SSTs. Separate experiments indicated that this intensity of photoactivating light decreased SST activity by a factor of ∼1.9(Heintz et al. 2022). The red trace shows the model prediction when SST activity within the model was decreased by this factor. Silencing of SSTs resulted in an increase in response and a shift to more depressing dynamics (bottom, RMSE = 0.23). **B.** As C but when PVs were optogenetically activated (top, by a factor of ∼2.5) or silenced (bottom, by a factor ∼2.0). Activation of PVs decreased the amplitude of PCs (top, RMSE = 0.19). Silencing of PVs resulted in an increase in response amplitude and shift to more sensitizing dynamics (bottom, RMSE = 0.29). **C**. Proportion of the initial “good” solutions to the model (chi-square < 12) that also provided good solutions incorporating the results of the optogenetic manipulations in A and B. Filtering of solutions on the basis of adding optogenetic conditions was carried out in series in the order 1-4. Applying filter 4 (PV inhibition) had little further effect on the number of good solutions i.e almost all good solutions incorporating manipulations 1-3 also predicted the effects of PV inhibition. **D**. Comparison of connection weights (mean ± SD) of all good solutions after applying optogenetic filters 1-3 and 1-4. All weights remain similar after the fourth filter and it is these averages that are presented in Figs. 6 and 7.

**Figure 6.**
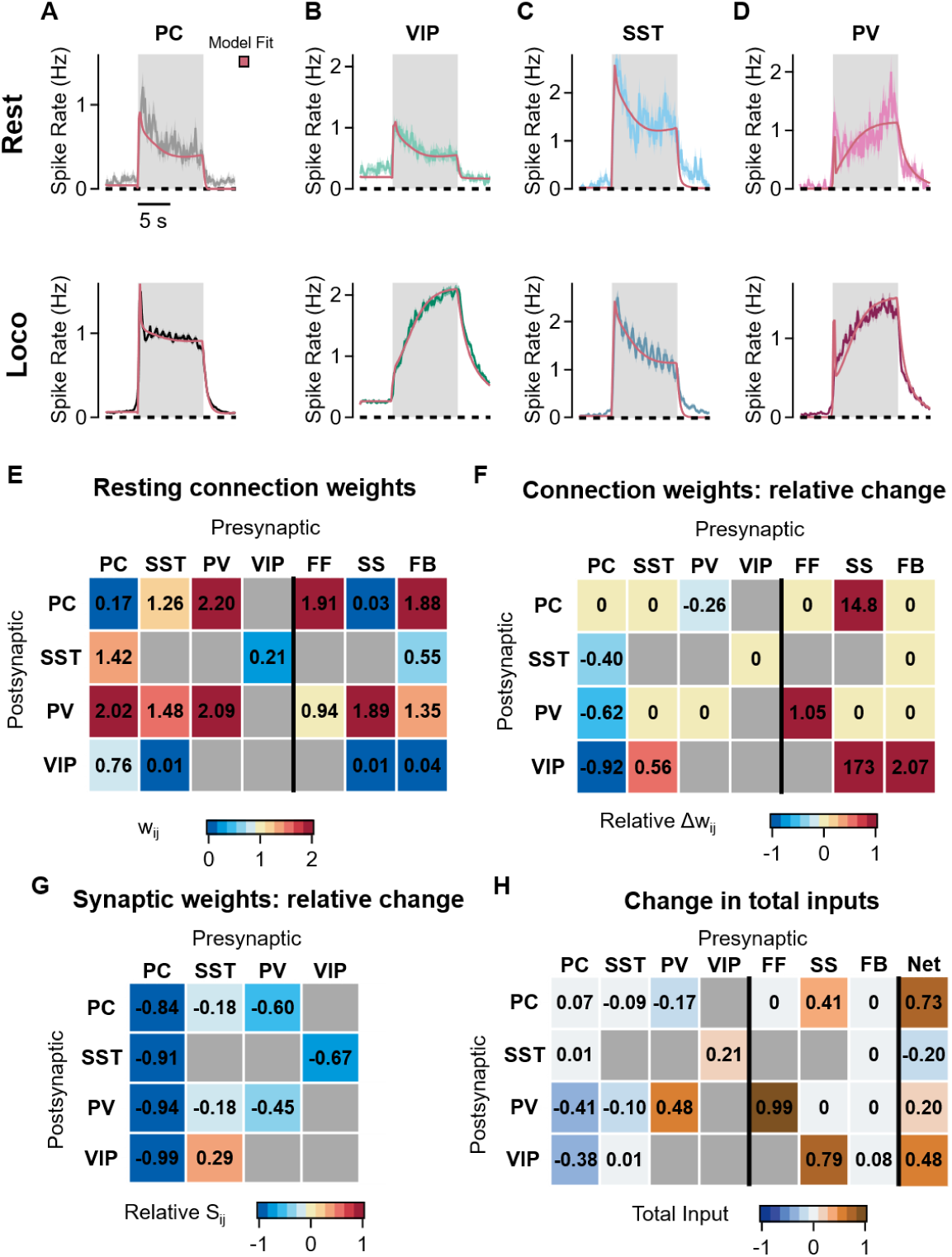
Locomotion modulates both electrical activity and the strength of synaptic connections. **A-D.** Top: Average firing rates of PCs (A, black), VIPs (B, green), SSTs (C, blue) and PVs (D, dark red) during stimulus presentation at rest and their corresponding fits from the average of solutions calculated by the model (light red, RMSE = 0.19). Note that these traces show the average activity of only partially paired recordings, where only a subset of cells are matched between rest and locomotion (rest n = 158 from 9 mice, locomotion n = 1437 neurons, from 14 mice). Bottom: average activity during locomotion in the same neuronal populations (RMSE = 0.11). **E.** Average resting connection weights from all solutions estimated by fitting the model to the activity of responsive neurons (Equation 2). **F.** Relative change between the time- independent connection weights inferred at rest and during locomotion (Equation 4). **G.** Relative change in synaptic weights based on estimated changes in connection weights **(F)** and measured changes in number of responsive cells (Supplementary Table 1; equation 3). Note that all excitatory and inhibitory synapses weaken except for synapses from SST to VIP which strengthen. **H.** The absolute change in the total synaptic input (locomotion-rest), calculated from the average activity (Supplementary Table 3) and the change in connection weight in F (equation 5). The last column is the net change taking into account the polarity of the synapse.

**Figure 7.**
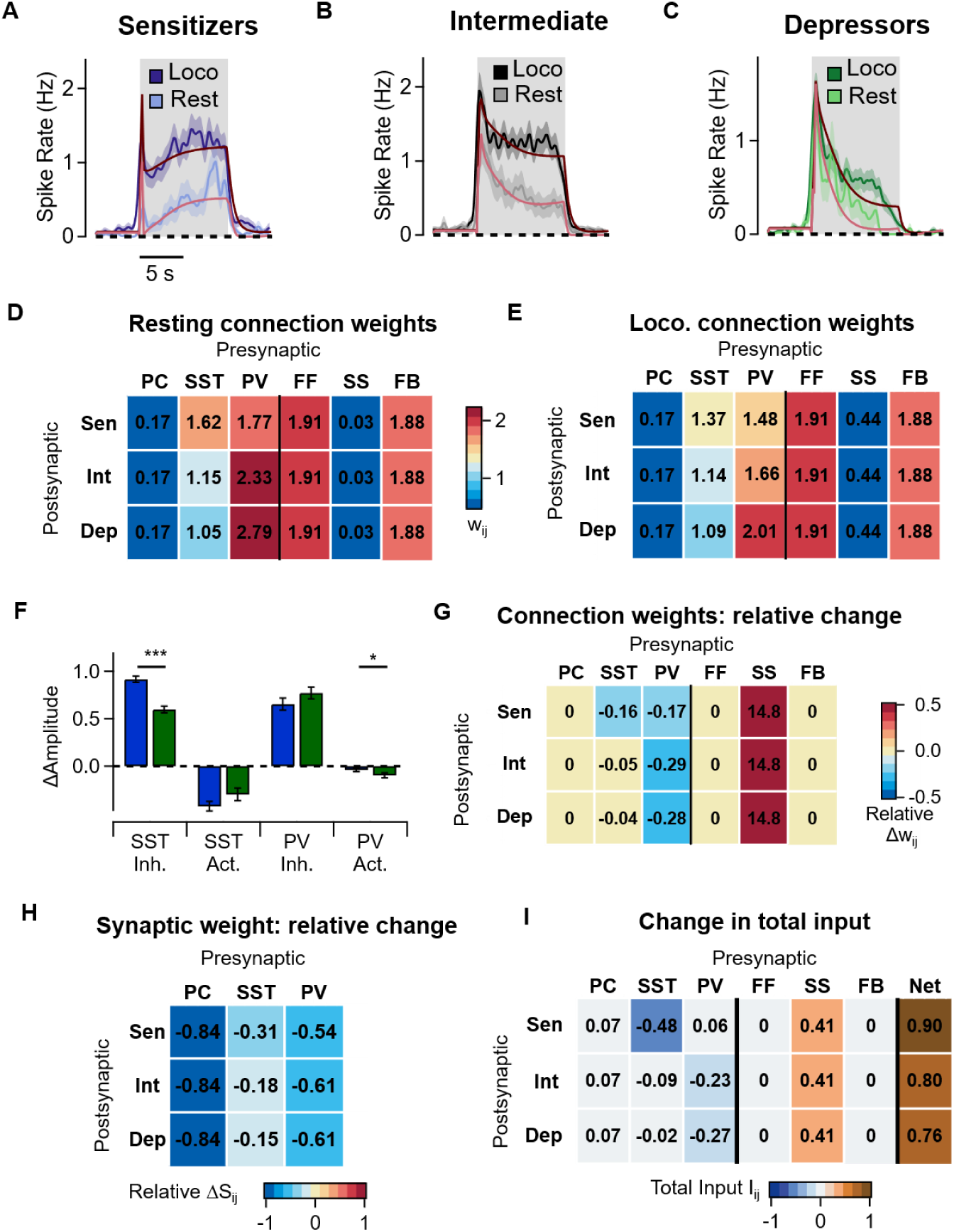
Differences in PV:SST input ratio determine the direction of PC adaptation and response amplification associated with locomotion. **A.** Activity of PCs from the tertile with lower AIs (sensitizers) at rest and during locomotion together with fits from the average of all good solutions calculated by the model (rest, RMSE = 0.22; loco, RMSE = 0.17). These are paired measurements from 158 neurons in 9 mice. During locomotion 7.5 ±1.2 more spikes were fired during the stimulus. **B-C.** As A, but for the intermediate and depressing tertiles, RMSEs were similar to sensitizers. During locomotion the stimulus generated 6.3 ± 0.9 more spikes in the intermediate group and 2.8 ± 0.4 in depressors. **D-E.** Heatmaps of the synaptic weights for each PC tertile at rest (D) and locomotion (E); sensitizers (Sen), intermediate (Int) and depressing (Dep). **F**. Change in PC response amplitude induced by optogenetic manipulation of SSTs and PVs in sensitizer (blue) and depressing (green) PCs. Note that sensitizers are preferentially modulated by SSTs and depressors by PVs as predicted by the model. (Inh: silencing with ArchT, Act: activation with ChrimsonR) (SST Inh: 1244 Cells from 5 mice; SST Act: 539 cells from 4 mice; PV Inh: 446 cells from 4 mice; PV Act: 180 cells from 3 mice. LMM: * p<0.05, *** p<0.001). **G**. Relative change in connection weights caused by locomotion for each subpopulation of PCs (calculated as equation 4). The key difference is the relative strength of PV and SST inputs. **H.** Relative change in synaptic weights caused by locomotion for each subpopulation of PCs. **I.** The absolute change in the total synaptic input (locomotion-rest) received by each subpopulation of PCs. Sensitizers receive less SST inhibition than the intermediate and depressing tertiles. The last column shows the *net* change in total input, taking into account the polarity of the synapse. There were no substantial changes between sensitizers and depressors (*U_norm_* = 0.4).

The first optogenetic manipulation was to over-activate SSTs by expressing the excitatory photoprotein ChrimsonR using a Cre-dependent viral vector. Simultaneously, responses were recorded in PCs using GCaMP6f under the CaMKII promoter. The intensity of the photo-activating light was adjusted to increase the activity of SST interneurons by an average factor of ∼2 in locomoting animals, as estimated in a separate series of experiments (Heintz et al. 2022). Fig. 5A (top) shows how the average PC response in the locomotion state was reduced compared to interleaved control trials.

Scaling the average SST activity within the model by a factor of 1.7 while leaving all other parameters constant provided an accurate description of the change in average PC activity in only 67% of the original solutions (Fig. 5C). In other words, although a broad region of parameter space could reproduce the control data, introducing this optogenetic manipulation eliminated roughly one-quarter of the solutions, thereby narrowing the admissible parameter space.

After this first filtering step, we sequentially reduced the number of good solutions by constraining to results of SST silencing, PV activation and PV silencing, in this order. Successive filters produced progressively smaller reductions in the number of solutions: SST silencing reduced them to 6% of the original, PV activation to 4% and PV silencing to 2% (Fig. 5C). Despite this further reduction, connection weights remained identical when applying the first three optogenetic constraints compared to applying the fourth (Fig. 5D). PV silencing therefore served as a validation step: it confirmed the region of parameter space defined by the three previous filters instead of further refining it.

### Locomotion increases PCs responses but weakens their local synaptic output

Behavioural modulation of connection weights *within* layer 2/3 have been very hard to quantify experimentally, making it unclear how far changes in internal state adjust cortical processing through changes in neuronal activity as opposed to modulation of synaptic weights (Kimura and Baughman 1997, Pronneke et al. 2015). We used the model constrained by the four optogenetic manipulations to infer changes in synaptic weights. First, we investigated the action of locomotion averaged across the whole PC population (Fig. 6) and then explored why the subset of PCs that sensitize is modulated more strongly than those that depress (Fig. 7).

The locomotion and rest datasets were fitted independently, yielding 1,007 and 11,657 final solutions, respectively. These solutions closely followed both the fast and slow phases of adaptation at rest and during locomotion, as exemplified by the response generated using the mean connection weights across the solution population (Fig. 6A- D, Supplementary Figure 4). Unsupervised clustering analysis identified a single cluster of solutions for locomotion and another for rest, indicating that these parameter sets form a continuum rather than falling into distinct categories. Moreover, examination of the distribution of each parameter revealed that they were mostly unimodal (Supplementary Fig. 4). The only exception was the PC->SST connection, which was described by two overlapping Gaussian components that were similarly reduced during locomotion. The absence of clearly separated modes suggests that the solutions did not form distinct families of network configurations. Together with the close fit produced using the mean connection weights, this supports their use as a representative population-level summary. The average connection weights (Equation 2, w_ij_) at rest are shown in the heatmaps in Fig. 6E and the relative changes with locomotion are summarized in Fig. 6F, calculated as:

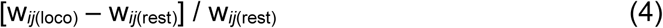

Where free parameters predicted by the model could be compared with published experiment, there were several points of agreement. For instance, the relative strength of direct local connections to PCs were quantitatively similar to those measured using electrophysiology (Jiang et al. 2015, Hage et al. 2022). There were, however, differences with other attempts to infer connection weights. Using our nomenclature, a recent modelling study (Kraynyukova et al. 2022) inferred a weight order in V1 at rest of *^w^PV*,*PC* ^< *w*^*PC*,*PC* ^< *w*^*PV*,*PV* ^< *w*^*PC*,*PV*, while our results infer *^w^PC*,*PC* ^< *w*^*PV*,*PC* ^< *w*^*PV*,*PV* ^<^ *w_PC_*_,*PV*_ (Fig. 6E). This difference may reflect our models incorporation of long-range modulatory inputs, which also allowed it to predict other experimental observations, including a locomotion-dependent amplification of the slowly modulated input to VIPs (Fu et al. 2014) and PCs (Jordan and Keller 2020). The incorporation of long-range modulatory inputs into the model were essential to accurately estimating connection weights within layer 2/3 because removing these inputs did not yield any solutions following the same quality criteria (Supplementary Fig. 5, see Methods).

The model indicates that locomotion increases the responses of PCs through a broad combination of circuit effects but of particular importance is a 18-60% reduction in the strength of inhibitory synapses made by SSTs and PVs (Equation 3, s_ij_; Fig. 6G). Strikingly, the model also predicts that the *total* weight of connections from PCs to all types of interneurons is decreased by 40–92% following locomotion (Fig. 6F). Taking into account the 5-fold *increase* in the number of responsive PCs during locomotion (Supplementary Table 1) this leads to an estimated 90% decrease in synaptic strength across all PC targets (Fig. 6G). The model indicates that the large increase in PC and PV activity in the locomoting state is associated with a wide-spread decrease in the strength of their local synapses. This mechanism would allow large changes in the gain of signals *leaving* V1 for higher visual areas but without overexcitation of PC targets *within* the local circuit. The possible cholinergic mechanisms underlying these changes are considered in the Discussion.

In contrast to the rest of the local network, the average SST->VIP connection weight *increased*, increased by 56% during locomotion (Fig. 6F). The number of responsive SST neurons increased by only 22% (Supplementary Table 1) indicating that the major cause was a *strengthening* of SST synapses (Fig. 6G). This differential pattern on excitatory synapses of PCs (suppression) and SSTs contacting VIPs (enhancement) could be consistent with known variations in the distribution of different muscarinic acetylcholine receptors (see Discussion). Switching to the active behavioural state also increased the strength of specific inputs to layer 2/3, most obviously the FF connections to PVs and the SS and FB connections to VIPs (Fig. 6F). VIPs were the most striking target, explaining their dramatic response increase (Fig. 3G).

Another way to summarize the effects of locomotion is to consider the change in the *total* synaptic input (both excitatory and inhibitory) that each neuron receives (Supplementary Table 3; Fig. 6H). The total synaptic input that neuron i receives from neuron j, I_ij_, is proportional to the activity of j and can be expressed as:

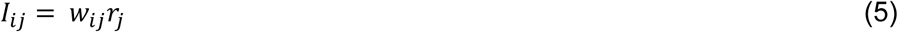

The last column in Fig. 6H shows this change, taking into account the input received from all connected populations averaged over the stimulus presentation. The net excitatory input driving the visual response is more pronounced in PCs and VIPs compared to PV and SST interneurons, consistent with the locomotion-associated increases in their responses shown in Fig. 3. The model predicted a modest net decrease in input to SSTs, although this was not accompanied by a detectable change in their average activity experimentally (Fig. 3I). The overarching picture is that locomotion adjusts the magnitude of visual responses by modulating both excitatory signals entering layer 2/3 and the strength of local synaptic connections, expanding what has been observed in a previous study (Dipoppa et al. 2018).

### Differential modulation of SST inputs onto sensitizing and depressing PCs

Why do some PCs sensitize while others depress? Optogenetic manipulations of interneurons indicate that the direction and amplitude of slow adaptation reflect the balance between PV and SST inputs (Heintz et al. 2022). PVs sensitize (Fig. 3H), so PCs in which these inputs dominate tend to depress, while SSTs depress (Fig. 3I), causing PCs in which these inputs dominate to sensitize. To make these ideas quantitative, we fit the model separately to the sensitizing, intermediate and depressing PC subpopulations (Fig. 7A-C). The simplest model that maintained a good-quality fit kept excitatory drive from PCs and all external inputs fixed and equal across the three PC types, restricting their variability to their local inhibitory connections (Fig. 7D,E). Similarly to the global model, the distribution of weights could be represented by the average but the whole distribution was considered during analysis (Supplementary Fig. 6). In qualitative agreement with the “balance model”, the direction of adaptation of a PC was dependent on the ratio of the PV and SST inputs, which increased progressively between the sensitizing, intermediate and depressing populations in both rest and locomotion (Fig. 7D,E).

We tested this prediction of the model by comparing the effect of optogenetic manipulations of SST and PV interneurons during locomotion, which provided qualitative support for the prediction that sensitizers are more strongly influenced by SSTs, whereas depressors are more strongly influenced by PVs (Fig. 7F). This balance also influenced the release from inhibition that different PCs undergo during locomotion, with sensitizers receiving less inhibition than depressors from SST interneurons (Fig. 7G; Sen PCs: - 0.16; Dep PCs: −0.04; U_norm_ = 0), while the inverse pattern is observed for PV connections (Fig. 7G; Sen PCs: −0.17; Dep PCs: −0.28; U_norm_ = 0.9). The adaptive properties of PCs reflected not only the relative contributions of local SST and PV inhibition but also changes in their adaptive dynamics and external inputs.

In principle, the differences in connectivity between sensitizing and depressing PCs could reflect synaptic strength (Equation 3) or total presynaptic inputs (Equation 5) received by the target cell. We found substantial differences in both the synaptic weight of SST and PV connecting to PCs (Fig. 7H), and in the total input these inhibitory cells convey to depressors and sensitizers (Fig. 7I). Again, the key difference was the PV:SST input ratio: sensitizers experienced a pronounced decrease in total SST inputs and a small increase in PV input (Fig. 7I). Notably, despite the larger locomotion-associated response increase in sensitizers, the model predicted comparable changes in net input to sensitizers and depressors (Fig. 7I). This dissociation is consistent with previous evidence that SST inhibition acts divisively rather than subtractively (Lee et al. 2012, Phillips and Hasenstaub 2016) (see Discussion). Overall, the model indicates that the enhanced responses observed in sensitizers is because they experience a greater reduction in SST inhibition during locomotion.

### Random inhibitory connections can account for opposite forms of adaptation

The model provided *quantitative* predictions of how the balance between PV and SST inputs to PCs varied to drive adaptation in opposing directions and to varying degrees. In sensitizing, intermediate and depressing PCs, the ratio of PV:SST inputs were 1.1, 2.0 and 2.7 at rest (Fig. 7D). During locomotion, these ratios were 1.1, 1.5 and 1.8 (Fig. 7E). How does this variability arise? One possibility is that there are *specific* connectivity rules for interneurons contacting PCs based on molecular and/or functional differences (Yoshimura et al. 2005, Runyan et al. 2010, Rossi et al. 2020) and that generates two general classes of microcircuit within layer 2/3, one causing PCs to depress and the other to sensitize. The distribution of AIs measured experimentally was, however, unimodal and did not provide statistical support for this idea (Fig. 1C). We therefore began by investigating the null hypothesis - random connectivity (Peters and Feldman 1976, Stepanyants and Chklovskii 2005, da Costa and Martin 2009, Rees et al. 2017, Schneider- Mizell et al. 2024). If variations in connectivity occur randomly, we can estimate the complete distribution of PV:SST input ratios from the known cell densities and average connection probabilities (Fig. 8A).

**Figure 8.**
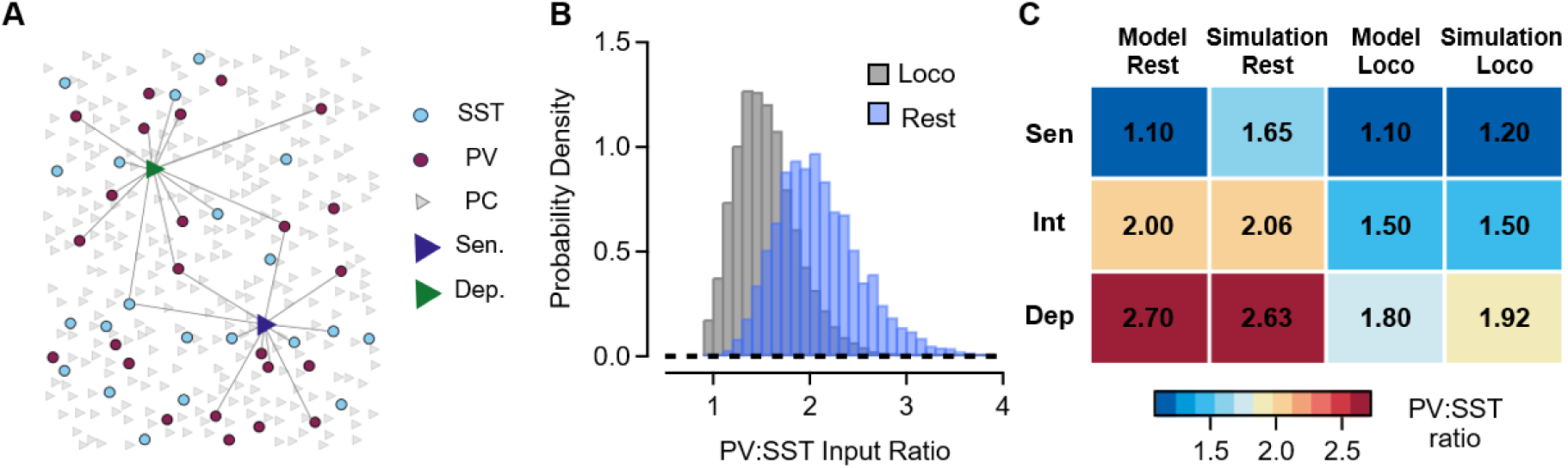
Random inhibitory connections to PCs can account for opposing forms of adaptation. **A.** Schematic of the simulation showing the relative densities of the four major types of neurons. Two PCs are highlighted with some of their connected interneurons (depressing, green; sensitizing, blue). The connection from a given interneuron to a PC was set by tossing a coin loaded according to the connection probability that has been measured by electrophysiology. **B.** At rest, the distribution of the PV:SST input ratios from a simulation of 100,000 PCs was skewed towards higher values; locomotion shifted the distribution to lower values. **C**. Comparison of the PV:SST input ratios calculated from the model (Fig. 7D and E) and from the simulation based on random connections. There is agreement to within 10% across a range of conditions, except for sensitizing neurons at rest.

Considerable efforts have been made to measure connection probabilities and synaptic strengths using electrophysiology. When recordings are made simultaneously from a PC and a nearby PV interneuron, the probability of a connection is ∼0.37 at distances <150 µm (Campagnola et al. 2022, Hage et al. 2022). The corresponding connection probability for SST interneurons is lower, at ∼0.27. These measurements can be combined with estimates of the numbers of interneurons within the typical connection distance of a PC (Methods). Across several studies, the densities of PCs, PV and SST interneurons have been estimated as 100,000, 7,856 and 6,771 cells/mm³, respectively (Rodarie et al. 2022), yielding estimates of approximately 41 connected PV and 26 connected SST interneurons per PC. Given that an individual PV synapse is approximately 1.55-fold stronger than an SST synapse (Jiang et al. 2015), these anatomical and electrophysiological measurements predict a PV:SST input ratio of ∼2.4 if all connected interneurons are activated during the visual response. However, at rest only 10.1/32.9 PV interneurons and 5.1/13.7 SST interneurons were visually responsive (Supplementary Table 1).

Simulations of PV and SST inputs onto each of 100,000 PCs were carried out by interrogating each interneuron within 150 μm of the PC and deciding if it was connected by flipping a coin loaded to an average probability of 0.37 for a PV and 0.27 for an SST interneuron. The resulting inputs were weighted by the corresponding visually responsive fractions of 0.307 and 0.372 (Fig. 8A and B; see Methods). A distance of 150 μm was used because almost all connections occur within this range (Jiang et al. 2015, Campagnola et al. 2022). At rest, the simulated distribution was skewed towards higher PV:SST ratios because of the higher density of PV interneurons and their higher connection probabilities (Fig. 8B). Dividing the distribution at rest into three equal groups according to PV:SST ratio yielded mean ratios of 1.65, 2.06 and 2.63. These values were similar to the ratios independently inferred by the circuit model for sensitizing, intermediate and depressing PCs (1.1, 2.0 and 2.7, respectively), with particularly close agreement for the intermediate and depressing populations (Fig. 8C). Thus, much of the heterogeneity in inhibitory input inferred from PC responses could arise simply from stochastic variation in the numbers of PV and SST interneurons contacting individual PCs, without requiring selective connectivity between inhibitory interneurons and different PC response classes.

We next asked whether the same framework could account for the change in inhibitory balance during locomotion. The fraction of visually responsive PV interneurons increased more strongly than that of SST interneurons during locomotion (18.4/32.9 compared with 6.2/13.7; Supplementary Table 1), which in isolation would shift the distribution towards greater PV dominance. However, the circuit model inferred a substantially larger locomotion-dependent reduction in PV→PC than SST→PC synaptic strength (∼60% versus ∼18%). Incorporating both effects shifted the simulated distribution towards lower PV:SST ratios (Fig. 8B). The lower, middle and upper thirds of the locomotion distribution had mean ratios of approximately 1.20, 1.50 and 1.92, closely matching the independently derived model estimates for sensitizing, intermediate and depressing PCs (1.1, 1.5 and 1.8, respectively; Fig. 8C). The model and simulations therefore agreed to within 10% across five of the six conditions. Thus, random anatomical connectivity can largely account for both the heterogeneity in PV:SST input across PCs and its reorganization during locomotion, reflecting state-dependent recruitment and modulation of inhibitory synaptic strength.

## Discussion

Adaptation is a fundamental aspect of sensory processing across cortical areas but it has not been clear how it is adjusted by changes in behavioural state that modulate these circuits. Here we have demonstrated that an active state acts on V1 to preferentially amplify responses in PCs that exhibit sensitizing adaptation over behavioural time- scales, with smaller effects on depressing PCs, and that this preferential amplification is most pronounced at lower running speeds (Figs. 1-3). A data-driven network model indicates that locomotion modulates both the external inputs arriving to V1 and local synaptic connections (Fig. 6). The differential modulation of the PC population reflects variations in their local inhibitory circuits: sensitizing PCs receive proportionally stronger SST input relative to PV input making them more susceptible to the locomotion-driven decrease in SST inhibitory strength (Fig. 7). More generally, these results build from the simple model of gain control through the VIP→SST→PC disinhibition pathway (Pi et al. 2013, Fu et al. 2014) by also considering the broader neuromodulatory actions of the cholinergic input to V1. The results, summarized in Fig. 9, indicate that locomotion increases PC responses through a combination of circuit effects, including an increase in external inputs and a decrease in local inhibition (Figs. 6 and 7).

**Figure 9.**
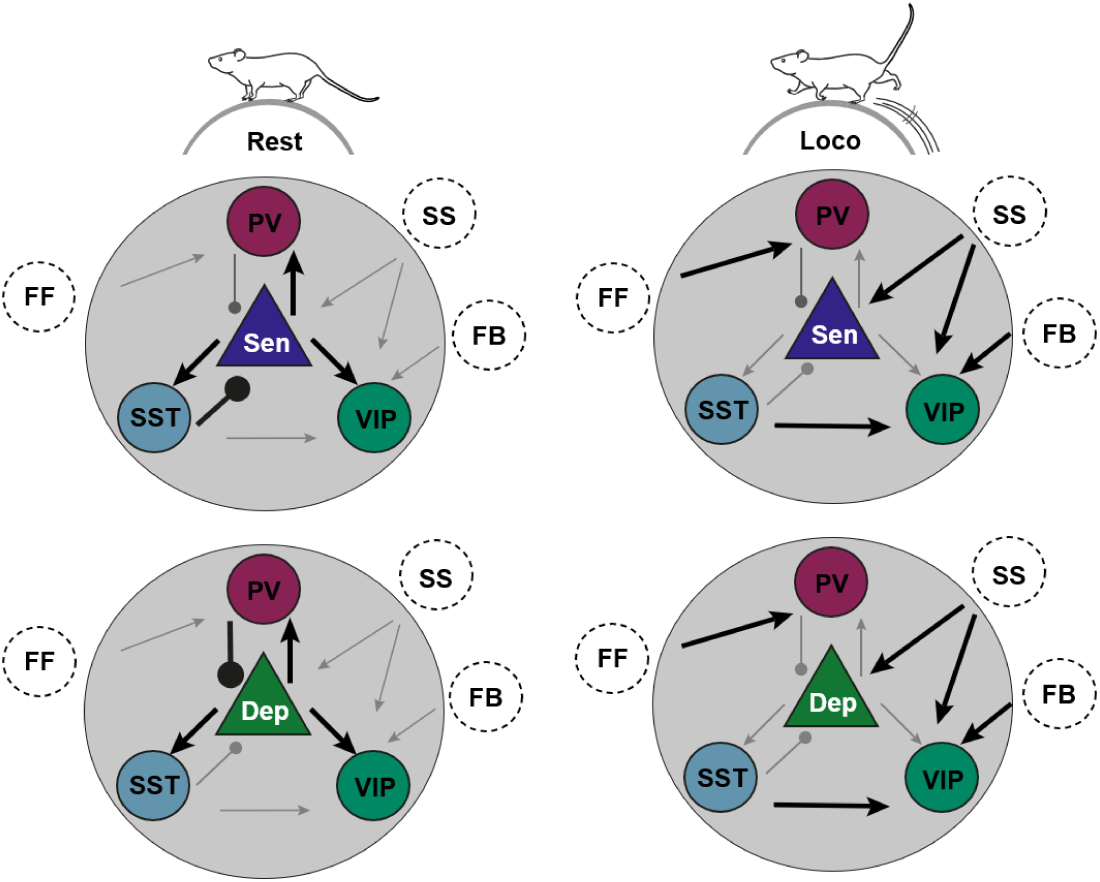
Summary of model-inferred circuit changes associated with locomotion in layer 2/3 of V1. Schematics summarise the circuit configurations inferred at rest (left) and during locomotion (right) for sensitizing (Sen; top) and depressing (Dep; bottom) PCs. Arrowheads and round terminals indicate excitatory and inhibitory connections, respectively; line thickness represents relative connection strength. Locomotion increases external drive to the local circuit while weakening excitatory connections from PCs to interneurons, as inferred from the population model in Figure 6. In contrast to the general weakening of local connections, SST→VIP inhibition increases during locomotion. Sensitizing and depressing PCs differ in the relative balance of their inhibitory inputs: sensitizing PCs receive proportionally stronger SST inhibition, whereas depressing PCs receive proportionally stronger PV inhibition, consistent with the subtype-specific model fits in Figure 7. Consequently, sensitizing PCs are more strongly affected by the locomotion-associated reduction in SST inhibition. The schematic provides a qualitative summary of the model results; connection strengths are not drawn to scale.

### Gating of the feedforward signal

The response amplification during locomotion was profound: the number of PCs responding to the stimulus increased by a factor of six (Supplementary Table 1) and the average amplitude of those responses by a factor of 2-3 (Figs. 1 and 3). About 70–80% of PCs in layer 2/3 send axon collaterals to higher visual areas (Glickfeld et al. 2013, Han et al. 2018) so these combined effects will strongly amplify the feedforward visual signal, at least to the high-contrast drifting grating stimulus used here. The local recurrent signals are, however, strongly damped by a wide-spread decrease in the strength of recurrent PC synapses subject to local modulation (Fig. 6). This decoupling of spiking activity and synaptic efficacy suggests a mechanism by which V1 can amplify its output to downstream targets, such as higher visual areas, without generating excessive excitation in the local recurrent network. Such a mechanism could be important for maintaining the stability of computations that rely on balanced excitation and inhibition within V1. We suggest that the suppression of PC output reflects a gating mechanism that allows feedforward communication to be selectively boosted while shielding local circuits from saturation. This prediction could be tested by performing paired electrophysiological recordings of PC→PC connections during state transitions, but direct measurements of changes in synaptic strength in the cortex of behaving mice are currently extremely challenging - most of the available evidence is inferred from circuit-level activity patterns and *in vitro* studies.

Acetylcholine release triggered by activity in the basal forebrain and prefrontal cortex acts across large areas through volume transmission (Ballinger et al. 2016, Lohani et al. 2022) but different muscarinic receptor types are expressed at specific locations, very often presynaptically, which provides spatiotemporally specific modulation(Munoz and Rudy 2014). The reduction in strength of local PC synapses during locomotion inferred from the model is in line with the known actions of acetylcholine on presynaptic M2 receptors, which reduce glutamate release at PC terminals (Hasselmo and Bower 1992, Mrzljak et al. 1993, Kimura and Baughman 1997). Many cortical areas are subject to modulation through the cholinergic input that they receive (Ballinger et al. 2016, Lohani et al. 2022), raising the possibility that signals from the basal forebrain also contribute to the increase in external input predicted by the model (Figure 6F).

### Locomotion shapes cortical dynamics through multiple pathways

In a previous, smaller dataset, we qualitatively examined the effects of locomotion across the PC population and found that sensitizing and depressing responses remained balanced, consistent with recruitment of the VIP→SST and SST→PV pathways (Heintz et al. 2022). Our expanded quantitative analysis and modelling now suggest that locomotion acts through changes in both external inputs and recurrent connection weights. Similar circuit changes were inferred by modelling the initial responses of V1 neurons to stimuli of different sizes (Dipoppa et al. 2018). That study also predicted increased external excitation and reduced recurrent excitation, resulting in weaker effective SST inhibition, but attributed the locomotion-dependent change in VIP responses to recurrent input from PCs rather than external drive. This difference may reflect the visual stimuli used. We used prolonged, high-contrast stimuli to provide a high signal-to-noise ratio and reliably evoke slow adaptation, whereas stimuli of different durations or contrasts may engage different patterns of locomotion-dependent modulation.

The first effect of locomotion we found was to enhance a slow sensitizing signal activated by the visual stimulus, the major target of which was VIP interneurons. The source of this signal might be the higher visual areas (Leinweber et al. 2017) and be gated by acetylcholine, before acting through VIPs to shape SST adaptation, with this final step supported by our optogenetic experiments (Fig. 4). Although the model predicted decreased net input to SSTs during locomotion, their average activity changed little. This apparent discrepancy may reflect opposing effects at different times: rapid nicotinic recruitment of VIPs could transiently supress SSTs, whereas slower metabotropic or recurrent mechanisms could subsequently restore their activity. Such time-dependent changes in connectivity are not captured by the time-independent weights in our model and may also be blurred by calcium-indicator kinetics.

The second major effect predicted by the model was an over 90% reduction in the strength of PC synapses across all targets (Fig. 6G), consistent with a range of evidence obtained *in vitro* demonstrating that M2 muscarinic receptors cause presynaptic inhibition (Hasselmo and Bower 1992, Gil et al. 1997, Kimura and Baughman 1997). A similar mechanism likely operates at PV, where release probability is reduced by M2 and M4 receptors (Xiang et al. 1998, Kruglikov and Rudy 2008, Urban-Ciecko and Barth 2016). However, the situation is different for SST interneurons. Our modelling shows that VIPs receive weak SST inputs at rest and that these synapses are *potentiated* during locomotion (Fig. 6E-G). This is likely to reflect a post-synaptic action of M1 receptors which have been shown to enhance postsynaptic GABA_A_ currents in visual and somatosensory cortices (Chen et al. 2015, Urban-Ciecko and Barth 2016). In contrast, SST inputs to PCs and PVs did not change during locomotion.

SSTs are a heterogenous class of inhibitory cells that target other cortical neurons in two distinct ways. The most numerous subtype, Martinotti cells, synapse on distal apical dendrites of PCs in layer 1 and show strong depressing adaptation, while Non-Martinotti cells generate more variable responses (Wang et al. 2002, Silberberg and Markram 2007, Adesnik et al. 2012). In line with this heterogeneity, the PC→SST connection showed a bimodal distribution at rest, with both components shifting similarly towards lower values during locomotion. We suggest that SSTs contacting VIP interneurons are predominantly Non-Martinotti cells because they are the only subtype known to be enhanced by acetylcholine through M1 and M3 receptors *in vitro* (Muñoz et al. 2017). It may also be that other neuromodulatory systems contribute to these mixed effects, including noradrenaline, serotonin and dopamine (Polack et al. 2013, Fu et al. 2014). The models might be refined by splitting SSTs into two populations once their responses are measured separately (Tremblay et al. 2016), but a more critical next step is to test the predicted changes in synaptic strength. As for PC→PC connections, the best way to test this would be to make paired electrophysiological recordings of identified SST→PC and SST→VIP connections during state transitions.

A key insight from our modelling is that the relative strength of PV and SST inputs predicts both the direction of PC adaptation *and* the degree of locomotion-induced response amplification (Fig. 7). Depressing PCs receive stronger PV input, consistent with their rapid and temporally precise inhibition (Atallah et al. 2012). In contrast, sensitizing PCs receive relatively stronger SST input which slowly depress to drive sensitization (Heintz et al. 2022). These differences in PV:SST ratio may also contribute to the response changes associated with locomotion. Although total SST input to PCs decreased only modestly in the population-average model (Fig. 6H), fitting the tertiles separately revealed a pronounced decrease in sensitizers but little change in intermediate or depressing PCs (Fig. 7I), explaining the weak effect in the pooled population. Thus, PCs already dominated by SST input underwent the greatest release from inhibition and the largest response enhancement. More generally, a broad weakening of inhibitory transmission would have a greater functional impact on PCs receiving stronger baseline input from that pathway, without requiring selective modulation of those connections.

### Modulation of SST→PC inputs during changes in brain state

An intriguing prediction of our model is that SST→PC synapses onto sensitizing PCs weaken whereas their PV→PC inputs change little during locomotion (Fig. 7I). As a result of the opposing effects of inhibition, the total net input does not differ substantially from that of depressor PCs. There has been controversy regarding the computational form of SST and PV inhibition onto PCs: under certain experimental conditions SST inhibition has been described as subtractive and PV divisive (Wilson et al. 2012), while different manipulations show the opposite (Lee et al. 2012, Phillips and Hasenstaub 2016). Although in our model we initially considered both types of inhibition subtractive, one possibility is that SST interneurons induce divisive inhibition during rest, such that their effect strengthens disproportionately with increasing inhibition. This may therefore account for the marked response increase we observed in sensitizer PCs, which receive stronger SST inhibition. In principle, this prediction could also be tested by targeted recordings of identified SST→PC connections during state transitions. This reduced model provided the simplest account of the data; however, because external inputs were not measured experimentally, their equality across PC subpopulations should not be interpreted as evidence that they are biologically identical.

Neural responses did not simply increase monotonically with running speed but rather reached its maximum at relatively low speeds, followed by a second, smaller maximum at higher speeds (Fig. 2). These maxima approximately coincided with speeds associated with walking and trotting, respectively (Bellardita and Kiehn 2015). Previous studies have reported a non-monotonic dependence of response gain and locomotion speed (Saleem et al. 2013), but our results suggest that this relationship may involve distinct locomotor regimes. One possibility is that changes in gait, and the associated differences in step dynamics or motor output, contribute to this pattern. Simultaneous measurements of gait and neuronal activity are, however, necessary to test this hypothesis.

What are the consequences of the enhanced responses of sensitizing PCs during locomotion? One possibility is that these changes maintain responsiveness to low contrast and transient stimuli in a visual environment that has become more dynamic with a strong component of visual flow. By selectively enhancing PCs that resist rapid depression, the circuit may preserve sensitivity to behaviourally relevant stimuli (Bennett et al. 2013, Dadarlat and Stryker 2017). From a computational perspective, the ability to differentially modulate adaptive subpopulations allows the cortex to tune its operating regime to changes in behaviour. Locomotion not only modulates the synaptic and gain mechanisms we focus on here but can also enhance neural representations of visual stimuli in V1 (Dadarlat and Stryker 2017). Our findings might, therefore, also be interpreted in terms of broader theories of predictive coding and attention (Rao and Ballard 1999): locomotion likely reflects an internal state of heightened predictive engagement with the environment and selective, speed-dependent amplification of sensitizing PCs could in principle support enhanced cortical representations of expected sensory features.

### Random local connectivity as a driver of functional heterogeneity

The use of a circuit model that incorporates measurements of activity from the three major classes of interneuron (Figs. 1-4) provided quantitative estimates of the way in which variations in local inhibitory circuits generate depressing and sensitizating modes of adaptation in PCs (Figs. 5-7). The balance between PV and SST inputs is a key parameter because these interneuron populations adapt in opposite directions to the visual stimulus (Fig. 2). Variations in this balance could be accounted for by two general statistics of the cortical connectome operating under a random connectivity rule (Fig. 8). First, interneurons occur at much lower density than PCs (Rodarie et al. 2022). Second, an individual interneuron only contacts a small fraction of nearby PCs. These results provide a stark example of the impact of Peters’ rule on cortical processing (Peters and Feldman 1976, Stepanyants and Chklovskii 2005, Braitenberg and Schüz 2013, Rees et al. 2017, Schneider-Mizell et al. 2024).

A random connectivity rule underlying variations in adaptation is wholly compatible with the idea that specific connectivity rules also operate within layer 2/3. It is clear, for instance, that SST interneurons target PCs while avoiding contact with each other (Pfeffer et al. 2013), while PCs of similar tuning are more likely to be connected (Ding et al. 2024). A recent connectomic study of mouse visual cortex compared the number of inhibitory synapses on the soma and dendrites of PCs (roughly corresponding to PV and SST inputs) and found a correlation coefficient of ∼0.5 in layer 2/3 (Schneider-Mizell et al. 2024) leading to the conclusion that the number of PV and SST inputs are determined by a combination of spatial overlap and more specific synaptic targetting. The present work complements such connectomic data by characterizing the functional impact of the differences in local inhibition that arise randomly. Such a “statistical summary of the connectome” (Rees et al. 2017) will be an important tool in understanding cortical function. We suggest that such a summary should include not only general circuit motifs but also the distributions that describe how these motifs vary (Fig. 8B).

There are large differences in the densities and connection probabilities of excitatory and inhibitory neurons in different areas of cortex, between layers within an area and even at different depths within a layer (Tremblay et al. 2016, Schneider-Mizell et al. 2024). For instance, the density of pyramidal cells in layer 2/3 of somatosensory cortex is about five times that in auditory cortex (Kim et al. 2017, Rodarie et al. 2022). There, the ratio of excitatory to inhibitory neurons is only 2:1 compared to ∼6:1 in auditory cortex. Even within layer 2/3 of visual cortex, the probability of a connection between an SST interneuron and pyramidal cell is lower in the more superficial zone (Hage et al. 2022). We are far from understanding the functional impact of these variations but this study indicates that the PV:SST input ratio may be a useful summary metric for understanding how inhibition alters PC function.

## Acknowledgements

This work was supported by BBSRC grant BB/X009386/1, a PhD scholarship to SD from the Leverhulme Trust (DS-2017-011) and a Researchers at Risk Fellowship to YK from the British Academy and Council for At-Risk Academics (RaR/100503).

## Methods

### Key resources table

| REAGENT or RESOURCE | SOURCE | IDENTIFIER |
| --- | --- | --- |
| Bacterial and virus strains |  |  |
| AAV1 | Addgene | 100834 |
| AAV1 | Charité | VCA-222a |
| AAV5 | Addgene | 28305 |
| AAV9 | Addgene | 100835, 62723 |
| Experimental models: Organisms/strains |  |  |
| VIP tm1(cre)Zjh/J | Jackson | #010908 |
| SST tm2.1(cre)Zjh/J | Jackson | #013044 |
| Pvalb tm1(cre)Arbr/J | Jackson | #008069 |
| Sst tm3.1(flpo)Zjh | Jackson | #028579 |
| Recombinant DNA |  |  |
| AAV1.CaMKII.GCaMP6f.WPRE.SV40 | Addgene | 100834 |
| AAV9.CAG.Flex.GCaMP6f.WPRE.SV40 | Addgene | 100835 |
| AAV1.EF1a-FLEX(frt)-GCaMP6f-WPRE | Addgene | 118273 |
| rAAV9/Syn.Flex.ChrimsonR.tdTomato | Addgene | 62723 |
| AAV5.CBA.Flex.ArchT.tdTomato.WPRE.SV40 | Addgene | 28305 |
| Software and algorithms |  |  |
| ScanImage | MBF Bioscience | ScanImage v.5 |
| PsychoPy | Peirce (Peirce 2007) | PsychoPy v. 1.82.02 |
| IgorPro | Wavemetrics | IgorPro v.8 |
| Suite2P | Pachitariu et al. (Pachitariu et al. 2016) | MATLAB version |
| MATLAB | MathWorks, Inc | MATLAB 2021a |
| MLSpike | Deneux et al.(Deneux et al. 2016) | MLSpike v.1.0 |
| V1 Model | Laboratory of Leon Lagnado | https://github.com/lagnadoLab/CortexModel |

### Experimental model

#### Animal preparation

All experimental procedures in mice were conducted according to the UK Animals Scientific Procedures Act (1986). These were performed at University of Sussex under personal and project licenses granted by the Home Office following review by the Animal Welfare and Ethics Review Board.

The animals were housed in polypropylene cages (21 cm × 35 cm) that contained nesting material, a shelter, a treadmill, a tunnel and wooden sticks. Mice were housed in groups of 2 - 3, except in some instances when reintegration into a group was unsuccessful after surgery. All animals were kept on an inverted 12-hour light-dark cycle. Results are reported from four VIP-Cre mice (VIP tm1(cre)Zjh/J Jackson #010908), fourteen SST-Cre (SST tm2.1(cre)Zjh/J, Jackson #013044), four SST-Flp (Sst tm3.1(flpo)Zjh; Jackson #028579) x VIP-Cre and ten PV-Cre (Pvalb tm1(cre)Arbr/J, Jackson #008069), all of them adult mice aged from P90 to P260 from either sex.

Mice were prepared for head-fixed multiphoton imaging through a surgical procedure in a stereotaxic frame under isoflurane anaesthesia. A titanium head plate was attached to the skull and a 3 mm craniotomy was drilled to expose the brain as described in detailed previously (Heintz et al. 2022). We then expressed AAV1.CaMKII.GCaMP6f.WPRE.SV40 to image calcium activity in pyramidal cells, AAV9.CAG.Flex.GCaMP6f.WPRE.SV40 to image interneurons in Cre lines and AAV1.EF1a-FLEX(frt)-GCaMP6f-WPRE to image SSTs in Flp lines. To excite interneurons optogenetically we used rAAV9/Syn.Flex.ChrimsonR.tdTomato and to inhibit them we used AAV5.CBA.Flex.ArchT.tdTomato.WPRE.SV40.

### Method details

#### Multiphoton imaging in vivo

During imaging sessions mice were free to run on a cylindrical treadmill placed under a two-photon microscope, with stimuli being presented on two monitors covering most of the visual field. Visual stimuli were sinusoidal gratings drifting upwards (100% contrast, 1 Hz, 10 s duration, spatial frequency 0.04 cpd, temporal frequency 1 Hz) generated using the Python library PsychoPy (Peirce 2007). The standard experimental protocol consisted of ten presentations of the stimulus with a 20 s interstimulus interval. We only analyzed responses to the first day of imaging in mice naïve to the stimulus to limit the introduction of longer-term changes in circuit function reflecting simple forms of learning such as habituation(Hinojosa et al., 2026).

When testing optogenetic manipulations these 10 trials were interleaved with 10 additional trials paired with illumination from an amber LED. The intensity of this LED was adjusted to activity of PCs to similar levels in different experiments. The two-photon microscope (Scientifica SP1) employed a 16x water immersion objective (0.8 NA; Nikon) and imaging was carried out at a depth of 150-300 μm below the surface of the brain.

Raw movies were registered and segmented into regions of interest (ROIs) using the Suite2P package running in MATLAB 2021a (Pachitariu et al. 2016) and further analysed using custom-written code in Igor Pro 8 (Wavemetrics). ΔF/F_0_ was calculated from the raw fluorescence trace using the mode of the entire trace as F_0_. For the control analysis in Supplementary Fig. 2E,F, ΔF/F responses were z-scored by subtracting the mean baseline ΔF/F and dividing by its standard deviation; the baseline period was defined as the interstimulus interval excluding the first 3 s after stimulus offset to allow signal decay. Here we only analyse results from the first imaging session in which the animal was exposed to the stimulus to avoid the effects of habituation observed over multiple sessions. Stimulus trials were only included for analysis if the mouse was running faster than 3 cm/s for more than 80% of the time. To calculate the response dynamics of PC, PV, SST and VIP populations we only included cells that were positively correlated with the stimulus (Excluded cells PCs: 30%; VIPs: 17%; PVs: 16%; SSTs: 50%). For each cell, the Pearson’s correlation coefficients was computed between its activity trace (ΔF/F) and a stimulus trace binarized as either stimulus off (0) or stimulus on (1) using the StatsLinearCorrelationTest Function in IgorPro (Wavemetrics). The threshold for significant positive or negative correlation was set at p < 0.05. Measurements of adaptive properties were confined to cells which were significantly positively correlated with the stimulus. The adaptive index (AI) was calculated from the GCaMP6f signal as AI = (R1 − R2)/(R1 + R2) where R1 is the average response over the period 0.5 - 2.5 s after stimulus onset and R2 is the average at 8-10 s. As a result, adding the same constant to R1 and R2 will make all AIs closer to 0 which is what we observe from rest to locomotion.

Running speed was measured with a rotary encoder and recorded through the imaging software Scanimage (Scanimage5; Vidrio Technologies). Trials were considered as locomotion when an animal ran at a minimum of 3 cm/s for at least 80% of the 10 s stimulus presentation, and were considered rest when remaining still (under 3 cm/s) during the same fraction of time. Trials with transitions from rest to locomotion and vice versa were rare and were excluded to ensure a stable behavioural state throughout the trial (See Supplementary Fig. 1 for an example trace).

#### Estimation of firing rates

To estimate firing rates from the recorded ΔF/F data we used the MLspike algorithm using initial parameters expected for GCaMP6f (Deneux et al. 2016). Briefly, this finds the optimal spike train to fit the calcium fluorescence trace (ΔF/F) taking into account estimates of baseline fluorescence level and neuronal parameters including the amplitude of the unitary Ca^2+^ transient generated by a spike (A) and the decay time-constant of that transient (τ). These model parameters were obtained by maximum- likelihood autocalibration of ΔF/F traces, allowing conversion of relative fluorescence changes into inferred spike counts and corresponding firing rates.

In general, we followed recommendations and demos in the article and on the MLspike GitHub page. We set the range for A to be between 1 and 50% and τ was set at values from 0.05 to 3 seconds. The parameter A_max_, which corresponds to the maximum amplitude of detected events, was 9, and the saturation parameter was set to 0.002. Finally, the Hill coefficient for Ca^2+^ binding to GCaMP6f was varied in the range from 1 to 3.75 with step sizes of 0.25.

The initial component should be interpreted cautiously because of the limited time-resolution of calcium imaging and because spike inference relies on assumptions that were not directly validated in our experiments. Nevertheless, the initial peak was also recovered using a different spike inference method: CASCADE. Calcium traces were analysed using CASCADE (Ref. Rupprecht et al., 2021) with the pretrained Global_EXC_6Hz_smoothing200ms model. CASCADE automatically matched traces to the available noise levels and combined predictions across five network ensembles. The resulting inferred spike probabilities were compared with the MLSpike results.

#### Model of the circuit in layer 2/3 of V1

We built a rate-based data-driven population model of the network circuit in the mouse visual cortex layer 2/3 (code and example data sets at https://github.com/lagnadoLab/CortexModel). The purpose of the model was to understand adaptive gain control during rest and locomoting states of four main neuron populations (PC, PV, SST, and VIP). Model parameters, both fixed and fitted, are listed in Supplementary Table 2.

The network was represented by the system of four mean-field first order ordinary differential equations (equation 1) (Wilson and Cowan 1972). The neuronal input-output function, *f*(*x*), was a rectified power law (Rubin et al. 2015)

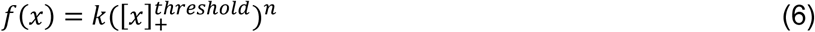

where *k* is a scaling coefficient which was set to 1. This is a rectified function with a lower threshold, 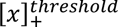, of zero to prevent negative values of firing rate and upper threshold of 25 to prevent run-away behaviour of the algorithm solving the network equations system. The exponent was set to n =2 but a linear activation function produced similar results during the fitting process, as observed by others (Kuchibhotla et al. 2017).

Connectivity between and within populations is represented by total connection weight *w_iii_* which is dependent on both the number of responsive cells in the presynaptic population and the strength of individual synapses. The number of presynaptic neurons responding to the stimulus varied between rest and locomotion so to assess changes in synaptic strength caused by neuromodulators the total connection weight was divided by the number of responsive presynaptic cells (Supplementary Table 1 and equations 2 and 3 in Results). The aim of fitting the model to the average activity traces of each neuronal population was to obtain connection weights, *w_iii_*. Weights were set to zero where published work indicated that connections are very weak or non-existent, as between SST interneurons (Pfeffer et al. 2013, Hage et al. 2022). The final connection weights matrix for connections within layer 2/3 was represented as:

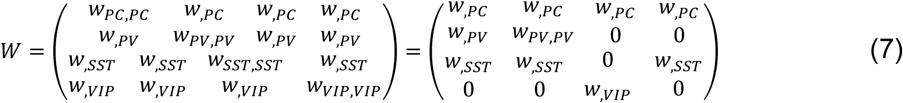

based on the reasonable assumption that connection probabilities are constant on the time-scale of minutes. Explain assumptions.

#### External inputs driving activity

Three types of external input drove activity in the circuit which we termed feedforward (FF), slow sensitization (SS) and feedback input (FB): the forms and origins of these are discussed in Results. These were represented in Equation 1 by the variable *In* that was specific for each input. The FF input was defined by the following system of equations:

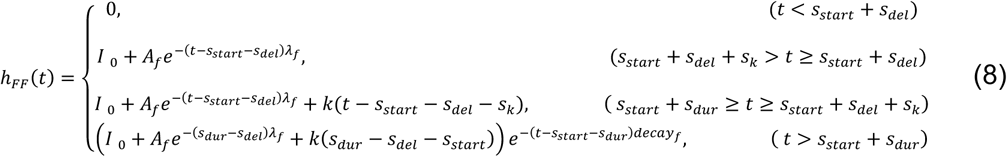

The initial short fast-depressing component had an amplitude *A_f_* and decayed exponentially with a decay rate *λ_f_*. The variables *s_start_*, *s_del_*, *s_dur_*, *s_k_* represent stimulus start, delay in the initiation of the response in a cell, the stimulus duration and an offset after the initial peak, respectively. This offset defined the initiation of a second slow increase in feedforward input that has been observed in response to drifting gratings lasting seconds (Makino and Komiyama 2015, Keller et al. 2020). At the end of the stimulus presentation, experimental traces decayed relatively slowly compared with the time-constant of the model neurons and this was accounted for by including an exponentially decaying component to the input with the argument *decay_f_*.

The temporal form of the feedforward input measured experimentally reflects the application of a spatio-temporal Gabor filter to a sinusoidal wave of drifting gratings (Gáspár et al. 2019). This oscillation was clear in the activity of individual PCs but less evident in population averages because neurons responded with different phases, depending on the location of their receptive fields relative to the stimulus. This 1 Hz oscillation was most clearly observed in the average response of the SST population (Fig. 5C, bottom) which have larger receptive fields and therefore display less phase shifting (Adesnik et al. 2012). We did not, therefore, attempt to model the oscillatory component of individual neuronal responses and concentrated on capturing the average activity across populations. The feedforward input (FF) was defined as a step of duration *s_dur_* with initial amplitude (*I*_0_ + *A_f_*) which then decayed to a steady state of *I*_0_ with time- constant 210 ms, defined by fitting the initial peak of the average PC response. The steady state was half the initial maximum with *A_f_* = *I*_0_. The linear increase is a simplified representation of the sensitizing behaviour observed experimentally in layer 4 activity (Makino and Komiyama 2015, Keller et al. 2020).

The slow sensitizing input was represented by the following system of equations:

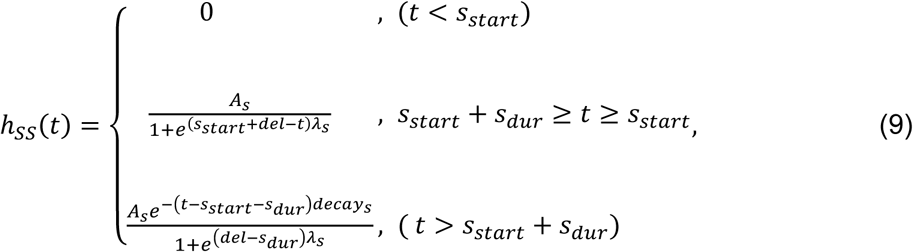

where *s_start_*, *s_dur_* represent the time of the stimulus start and the stimulus duration, respectively. The slow sensitizing (SS) component is represented by a sigmoid function with an amplitude *A_s_*, growth rate *λ_s_*, and centre of sigmoid shifted to a *del* value from the stimulus starting time. Similarly to FF, the decay phase after the stimulus end was accounted for by the argument *decay_s_*.

Finally, feedback input (FB) was represented by a step function:

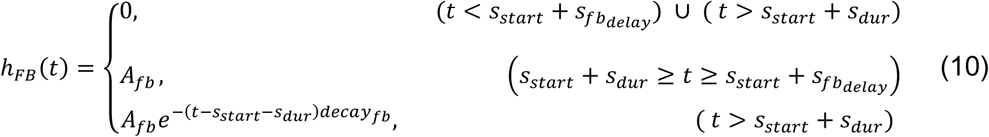

where *s_start_*, *s_fb_*__*delay*_, *s_dur_* represent time of the stimulus start, delay in the initiation of the response in a cell and the stimulus duration, respectively. The amplitude of the stimulus is defined by *A_fb_*. Similarly to FF and SS, the decay phase after stimulus end was accounted for by the argument *decay_fb_*. The amplitudes of A_fb_, A_s_ and I_o_ were fixed around 1 Hz based on the average response of PCs during locomoting state (Fig. 2).

#### Fitting the model

To obtain values of connection weights we fit the model simultaneously to the average activity traces of the four neuronal populations using the *LMFIT* library in Python(Newville et al. 2014) by minimization of reduced chi-square parameter *χ*^2^:

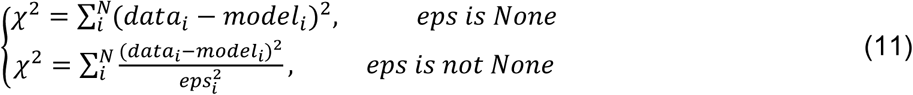

Where (*data_i_* − *model_i_*) represent an array of residuals between model and data points to be fitted, *N* – is the number of datapoints, and *eps_i_* is the uncertainty of the data points. The model was fitted to all time points of the average response traces, with differential weighting of specific time windows to capture both the initial and sustained components of the response.

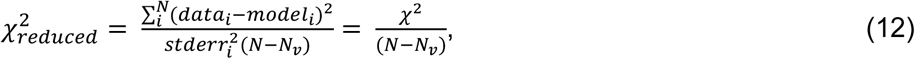

Where *N* is the number of data points and *N_v_* is the number of variable parameters. For the unrestricted four-population fit, there were *N* = 488 data points (122 in each cell) and *N_v_* = 27 varied parameters; in reduced models, *N_v_* decreased according to the number of parameters fixed. We set the threshold reduced chi-square value to be 12 for locomotion and 3 for rest, given their different variabilities. An ideal “good fit” would have a value of reduced chi-square around 1, as far as uncertainty is provided (Newville et al. 2014). We used standard errors of the mean (S.E.M) to weight the uncertainty of locomotion and rest data (eps). Additionally, to have a measure of absolute fit error independent of the data uncertainty and number of varied parameters we calculated root mean square error (RMSE) for each fit:

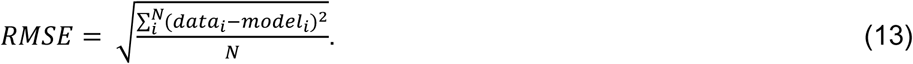

#### Parameter Search Strategy

Due to the high dimensionality of the parameter space, multiple parameter sets often produced similar 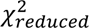 values, indicating that networks with different connectivity profiles could generate similar activity patterns, which in turn made the results sensitive to the initial guess of parameters. While global optimization algorithms such as genetic algorithm and dual annealing were tested, they did not yield satisfactory performance for this model.

Instead, having access to the Artemis high performance computing cluster at Sussex University, we combined Sobol sampling (Sobol 1976) with local optimization methods. Sobol sampling provides uniform coverage of high-dimensional parameter space. We generated 100,000 sets of evenly distributed initial parameters and each Sobol-sampled parameter set was subsequently refined using local optimization. We tested *Levenberg-Marquardt, Nelder-Mead, Powell* and *L-BFGS-B* methods and noticed that *Nelder-Mead and Powell* performed across a broad parameter range, whereas *Levenberg-Marquardt* and *L-BFGS-B methods* were more effective for final fine-tuning. This two-stage procedure (global Sobol sampling of the initial parameters followed by local optimization) allowed us to systematically explore parameter space and quantify the variability of estimated connection weights.

For the locomotion condition, 37 globally sampled solutions met the criterion for a good fit (reduced 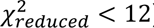). To generate a larger solution population for subsequent filtering, we repeated the procedure using regional Sobol sampling. We generated 80,000 additional initial parameter sets, restricting the starting value of each connection weight to the mean ± 2 SD of the 37 good global solutions, and then applied local optimisation without changing the fitting criterion. This produced 40,000 good solutions whose connection-weight means and SDs were similar to those of the initial solution population.

To reduce the number of free parameters and attempt to fit a simpler model, we measured the magnitude of the difference between rest and locomotion for each parameter. We quantified differences using the normalized Mann–Whitney statistic (See below), equivalent to the Vargha–Delaney *A* effect size (Vargha and Delaney 2000). Values above 0.7 or below 0.3 were considered large effects (Supplementary Fig. 3). As a complementary measurement to further reduce the number of free parameters, we performed principal component analysis on the difference between rest and locomotion solutions. As these were not paired, we sampled randomly both populations and calculated the difference of 1.000 pairs. We then calculated the variance weighted contribution of each weight to PC1 and fixed weights with borderline effect sizes and PC1 contribution below 0.5%.

An important additional constraint was provided by requiring the model to fit the results of optogenetic manipulations of the PV and SST populations (Fig. 5), leading to a reduced number of sets of possible synaptic weights. In these experiments, the intensity of the LED was set to increase or decrease PV or SST activity by a known factor between 1.6 and 2.5, as measured empirically in other parallel experiments measuring GCaMP6f responses in these interneurons (Heintz et al. 2022). The average PV or SST activity trace was then scaled by a similar factor for comparing the model to observed PC activity, followed by RMSE calculation for each comparison in optogenetic tests. We had a total of 4 conditions: SST activation, SST inhibition, PV activation and PV inhibition which we used for sequentially filtering the original group of solutions in 4 filtering steps. The threshold for a good opto fitting in each step was 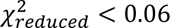 and 1.5 < OptoParam < 4. This approach allowed us to reduce the set of synaptic configurations, yielding parameter sets that captured both the average circuit dynamics during locomotion and the network’s response to optogenetic perturbations.

For the resting state fits, we repeated the Sobol sampling and local fitting procedure using the same initial steps excluding optogenetic filtering, which allowed us to estimate mean parameter values and their variability. We obtained 17 solutions through global sampling that met the fitting criterion (reduced 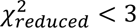), and 11,657 good solutions were subsequently produced following regional sampling and local optimisation. Optogenetic filtering was not feasible due to the noisier nature of these traces.

Alternative versions of the model were tested by setting the connection weights of the tested connections to zero, this was performed with the SS and FB inputs. Notably, all the resulting solutions had chi-squared values above threshold indicating that slow sensitizing and feedback inputs are essential to explain circuit dynamics on these time- scales (Supplementary Fig. 5).

To assess whether model parameters segregated into distinct groups, we applied unsupervised clustering using both K-means and DBSCAN algorithms. For K-means, we used the Elbow method and Silhouette analysis to estimate the optimal number of clusters. We further applied DBSCAN, estimating the neighborhood parameter (eps) from the 5th nearest neighbor distance using the KneeLocator method.

### Fitting the model to subpopulations of pyramidal cells

The whole PC population was split into tertiles (sensitizers, intermediate cells and depressors) based on the distribution of adaptive index (Fig. 1). The average response of each subpopulation was then fit separately for each of the good solutions from the locomoting and resting states using the two-stage fitting procedure explained above to assess changes in connectivity that describe the adaptive properties of each (Fig. 7). Three simplifying assumptions were made, based on what we ignore from the network. First, that each subpopulation of PC receives the same shape of inputs as the average PC across the population. Second, only the synaptic weight of those connections directly targeting PCs were left free to vary while the other connections within the circuit were fixed. This was done because we ignore if and how connectivity within the subpopulations of interneurons that target different PC type varies. For similar reasons, our third restriction was keeping external inputs and PC inputs fixed across PC types, because keeping them free did not improve the fitting considerably.

### Simulation of PV:SST input ratio based on random connectivity

Because the density of PC cells is higher than PV and SST, and because probability of connection from PV to PC and from SST to PC is lower than 1 there should be a broad distribution of the number of connected interneurons to PC and therefore distribution of proportional inhibition of PC cells by PV and SST. To test an assumption that randomness in connectivity between PV to PC and SST to PC cells can describe different polarities of slow adaptation observed in this work, we simulated 100000 PC cells and analysed a distribution of ratio of PV=>PC inhibition to SST=>PC inhibition. To be unbiased to the model results, we gathered all required information from the literature. Probabilities of connections from interneurons to pyramidal cells were set to *p_PV_* = 0.375 and *p_SST_* = 0.275 based on experimental data (Jiang et al. 2015, Campagnola et al. 2022, Hage et al. 2022). The average number PV and SST interneurons within 150 μm of each PC was 111 and 95, respectively, simply calculated from the measured densities of PCs (100,000 cells/mm^3),^ PVs (7856 cells/mm^3^) and SSTs (6771 cells/mm^3^) in layer 2/3 (Rodarie et al.). The strength of an individual synaptic connection was based on measurements of the unitary post-synaptic potential of *s_PV_* = 0.48 mV and *s_SST_* = 0.31 mV (Jiang et al. 2015).

Simulation for each pyramidal cell was performed in Python by drawing samples from a binomial distribution, represented as *numpy*. *random*. *binomial*(*n*, *p*, *size*) in numpy library, where *n* = 1 corresponds to number of trials, *p* - probability of connection, *size* - number of inhibitory cells. This is equivalent to tossing a coin with weights of p (connection) and 1-p (no connection) on either side. The coin is tossed 1 time for each interneuron around the picked pyramidal cell. The total ratio of PV:SST inhibition of PC cells is:

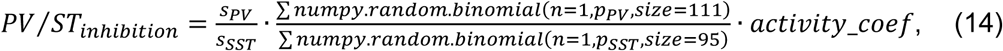

where *s_PV_*, *s_SST_* are average strength of synapses and *activity*_*coef* = 1.25 is the ratio of proportion of active PV and SST cells (0.81 and 0.65, respectively). The simulated distribution of PV:SST input ratios was skewed reflecting the higher density and connection probabilities of PV interneurons compared to SSTs (Fig. 8B).

### Statistical analysis

Because fitted model solutions do not constitute independent experimental replicates, conventional significance tests were not applied. Differences between solution distributions were instead quantified using the normalised Mann–Whitney U statistic, equivalent to the Vargha–Delaney effect size (Vargha and Delaney 2000):

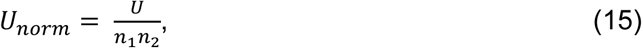

where U is the Mann–Whitney U statistic and *n*_1_, *n*_2_ are the sample sizes of the two groups. Rest and locomotion solutions were fitted independently and therefore had no natural one-to-one correspondence. To compare locomotion-induced changes between sensitizing and depressing PCs, we used Monte Carlo sampling to generate matched pairs of rest and locomotion solutions for each PC group and calculated the difference within each pair. The resulting distributions of locomotion-induced changes were then compared using *U_norm_*. A substantial increase in the direction Rest>Loco or Sen>Dep was considered when 1 > *U_norm_* > 0.7, a meaningful decrease in the direction Rest<Loco or Sen<Dep required 0.3 > *U_norm_* > 0, and 0.7 > *U_norm_* > 0.3 was considered as no change.

For the statistical tests not directly related with the model fitting we used the analytical software Igor Pro 8 and/or SPSS (’Statistical Package for the Social Sciences’, IBM). To determine if the mean of one group was significantly different from another group we used a linear mixed modelling (LMM) (Judd et al. 2017). Locomotion state, adaptive type, and their interaction were included as fixed effects, with mouse and cell identity as random effects. We tested differences among adaptive types separately within each behavioural state, differences between states separately within each adaptive type, and finally the interaction between behavioural state and adaptive type which indicates whether the locomotion-related increase in a given variable differed between sensitizers and depressors. Bonferroni correction was applied for multiple comparisons. All values are given as mean ± standard error of the mean (SEM) or mean ± standard deviation (SD) as indicated in each figure legend.

## Data and code availability

All code and data reported in this paper will be shared upon request. Code and example data sets are available at https://github.com/lagnadoLab/CortexModel.

## Supplementary Information

**Supplementary Figure 1.**
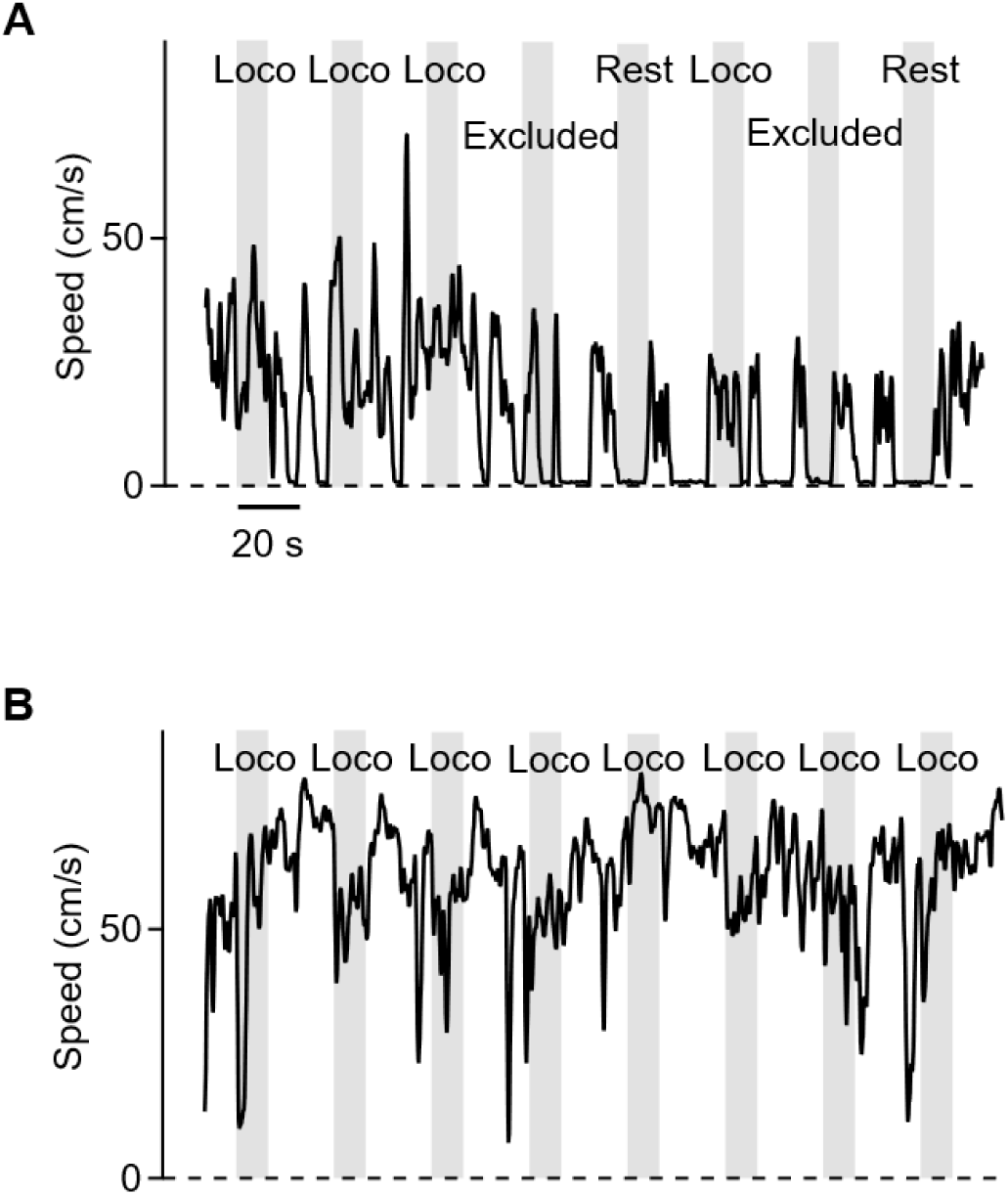
Example running-speed traces and trial classification. **A.** Example traces of running-speed from eight trials of the visual stimulus (grey bars). Trials were classified as locomotion when running speed remained above 3 cm/s for more than 80% of the trial. Trials were classified as rest when running speed remained below 3 cm/s for more than 80% of the trial. Trials not meeting either criterion were classified as mixed and excluded (see Methods). **B.** As in A, for a mouse that ran faster and more continuously.

**Supplementary Figure 2.**
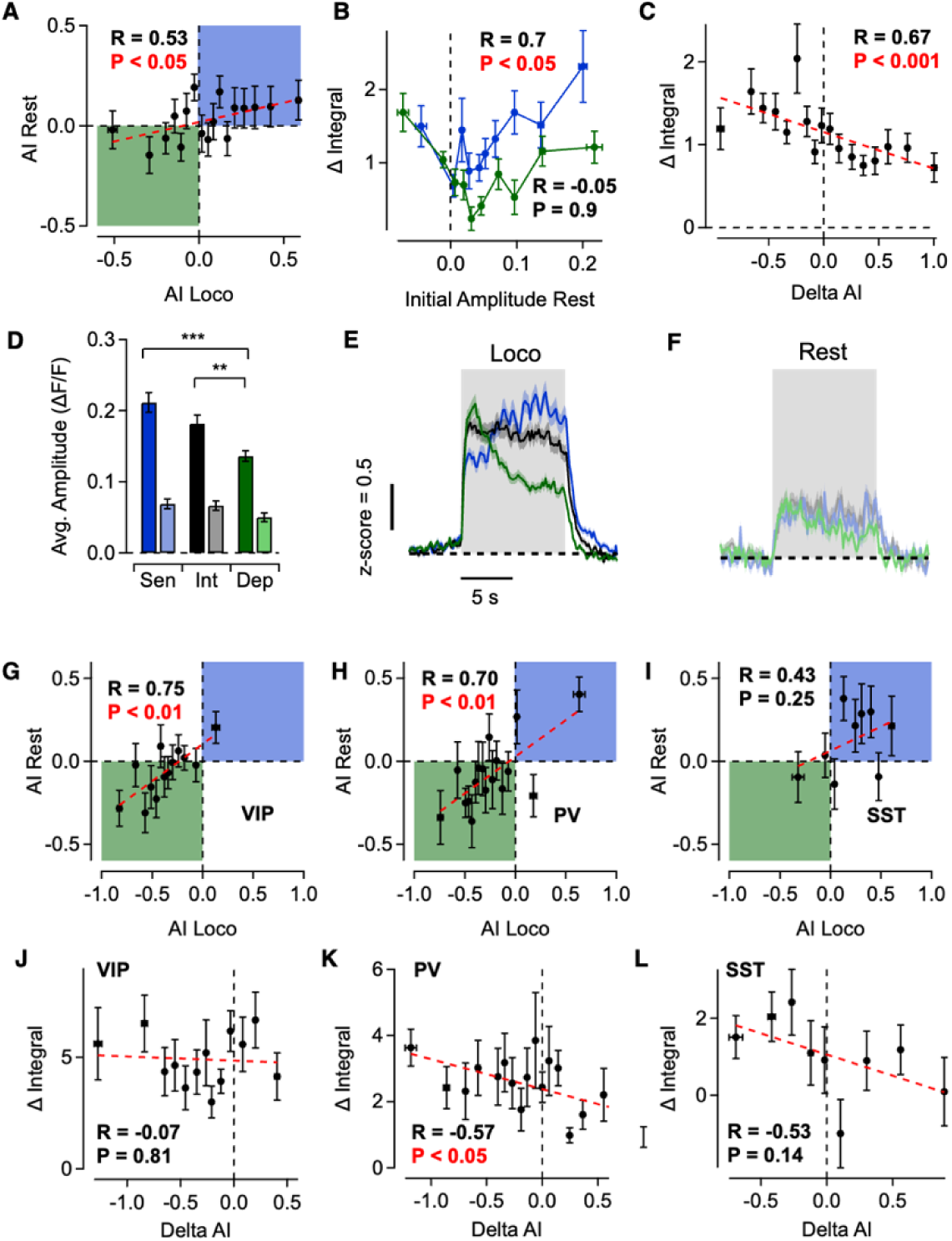
Locomotion-dependent changes in adaptive properties of PCs. **A.** Relationship between PC adaptive index (AI) at rest and during locomotion. AI values were positively correlated between behavioural states, indicating partial preservation of adaptive properties. Shaded regions indicate sensitizing (green; AI < 0) and depressing (blue; AI > 0) responses in both states. **B.** Locomotion-induced increase in response integral (ΔIntegral) as a function of initial amplitude at rest. Sensitizers increase in response integral is correlated with initial amplitude, but this is not observed in depressors. **C.** ΔIntegral as a function of the locomotion-induced change in adaptive index (ΔAI) in PCs. Cells that shifted towards sensitization during locomotion exhibited larger increases in response integral. **D.** Average response amplitude of sensitizing (Sen), intermediate (Int) and depressing (Dep) PCs at rest and during locomotion. Locomotion preferentially increased the response amplitude of sensitizing PCs similarly to ΔIntegral. **E–F.** Average z-scored responses of sensitizing (blue), intermediate (black) and depressing (green) PCs during locomotion (E) and at rest (F), showing a larger response magnification in sensitizers with similar baseline activities across populations and behavioural states. **G–I.** Relationship between AI at rest and during locomotion in VIP (G), PV (H) and SST (I) interneurons. AI was positively correlated between behavioural states in VIP and PV interneurons but not in SST interneurons. **J–L.** ΔIntegral as a function of ΔAI in VIP (J), PV (K) and SST (L) interneurons. A shift towards sensitization was associated with a larger locomotion-induced increase in response integral in PV interneurons, but no correlation was detected in VIP or SST interneurons. For scatter plots, cells were ordered according to the variable shown on the x-axis and grouped into bins of 30 cells for PCs and 15 cells for interneurons; correlations were calculated across the mean values of these bins.

**Supplementary Figure 3.**
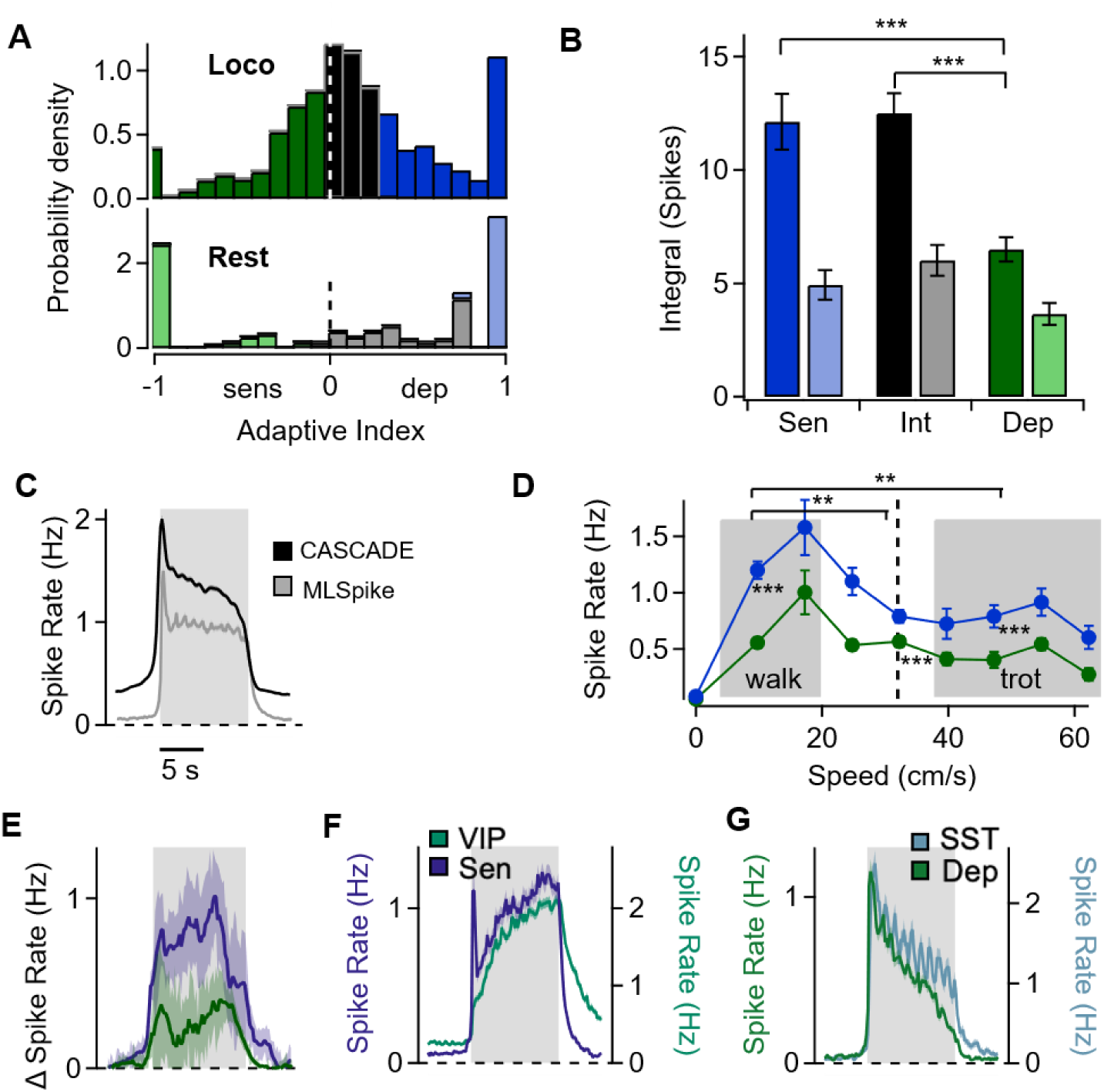
Locomotion preferentially increases the inferred firing rates of sensitizing PCs. **A.** Distributions of AIs calculated from inferred spike rates in the PC population during locomotion (top) and rest (bottom) in sensitizers (blue), intermediates (grey) and depressors (green). **B.** Response integral of sensitizing (Sen; blue), intermediate (Int; black) and depressing (Dep; green) PCs during locomotion (dark bars) and at rest (light bars). Locomotion preferentially increased the response integral of sensitizing PCs. **C.** Average inferred PC firing rate during locomotion calculated with CASCADE(black)(Rupprecht et al. 2021). This spike inference algorithm revealed a similar initial peak than MLSpike(grey). **D.** Locomotion-induced increase in response amplitude (ΔF/F) as a function of running speed in sensitizing (blue) and depressing (green) PCs. Cells were grouped in 7.5 cm/s speed intervals. Shaded regions indicate the complete bins included in the walking and trotting associated speed ranges, which were those whose interval overlapped the published gait-speed distributions(Bellardita and Kiehn 2015). The two ranges were compared with each other and with the intermediate-speed bin (dashed line); significance is indicated above the graph. Sensitizers and depressors were also compared within each speed range, with significance indicated adjacent to the corresponding range. **E.** Time course of the locomotion-induced change in firing rate in sensitizing and depressing PCs. **F.** Average inferred firing rates of sensitizing PCs (purple) and VIP interneurons (turquoise) during locomotion. The two populations exhibited similar sensitizing dynamics. **G.** Average inferred firing rates of depressing PCs (green) and SST interneurons (light blue) during locomotion. The two populations exhibited similar depressing dynamics. Grey shading indicates stimulus presentation. Data are mean ± SEM. **p** < 0.01, ***p*** < 0.001; linear mixed models.

**Supplementary Figure 4.**
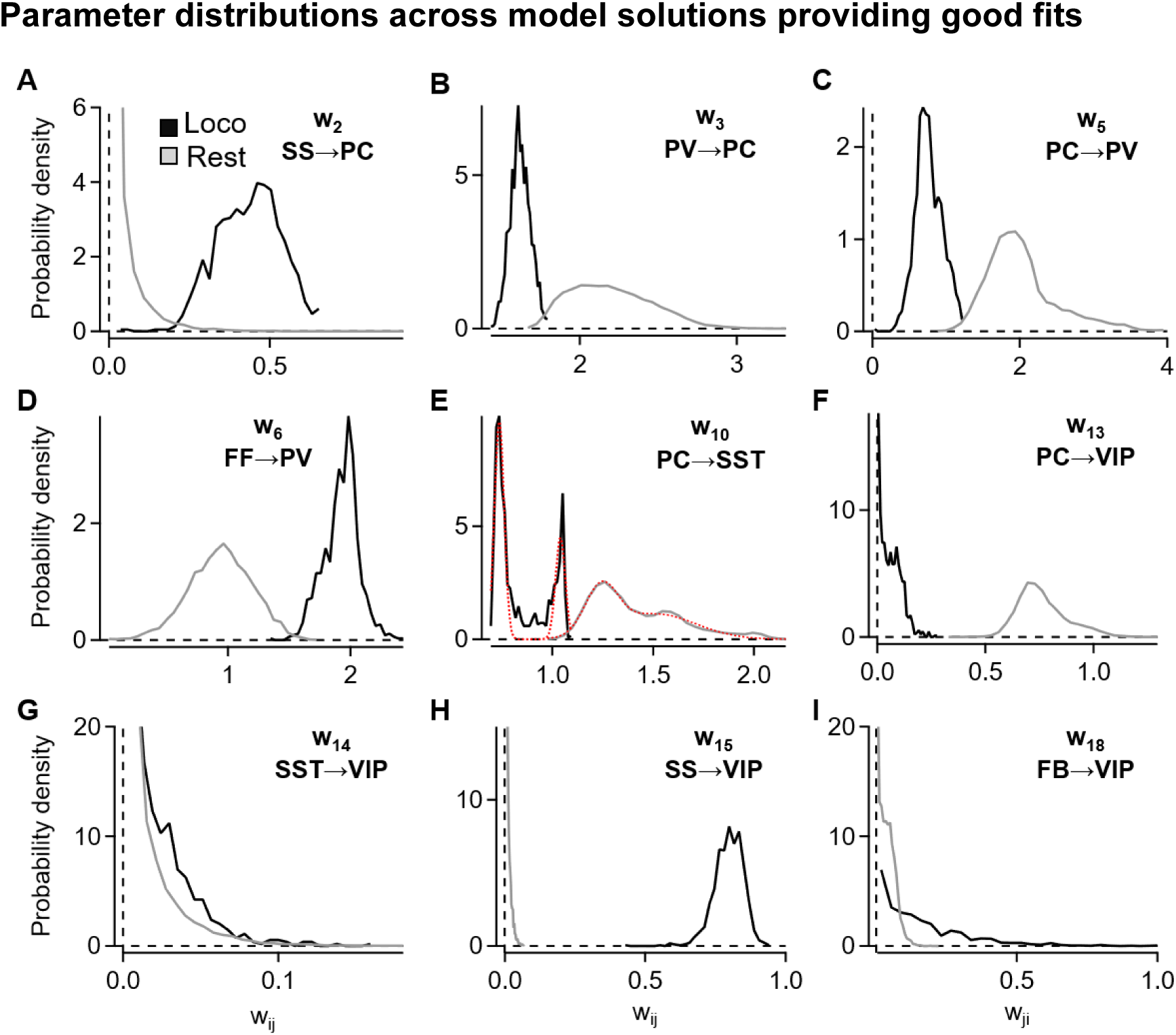
Parameter distributions across solutions providing good fits. Probability-density distributions of connection weights *w_iii_* across the accepted model solutions at rest (grey) and during locomotion (black). **A–I**. Distributions are for SS→PC (A), PV→PC (B), PC→PV (C), FF→PV (D), PC→SST (E), PC→VIP (F), SST→VIP (G), SS→VIP (H), and FB→VIP (I). Red dotted lines in E show fits with the sum of two Gaussian distributions (Rest: Peak 1 x_0_ =1.2±0.004, Peak 2 x_0_ =1.5±0.03; Loco: Peak 1 x_0_ =0.7±0.002, Peak 2 x_0_ =1.04±0.005), revealing a bimodal distribution of PC→SST weights in both conditions. Both components underwent a similar shift with locomotion (peak 1: 0.50; peak 2: 0.47), suggesting that the two components were similarly affected by locomotion.

**Supplementary Figure 5.**
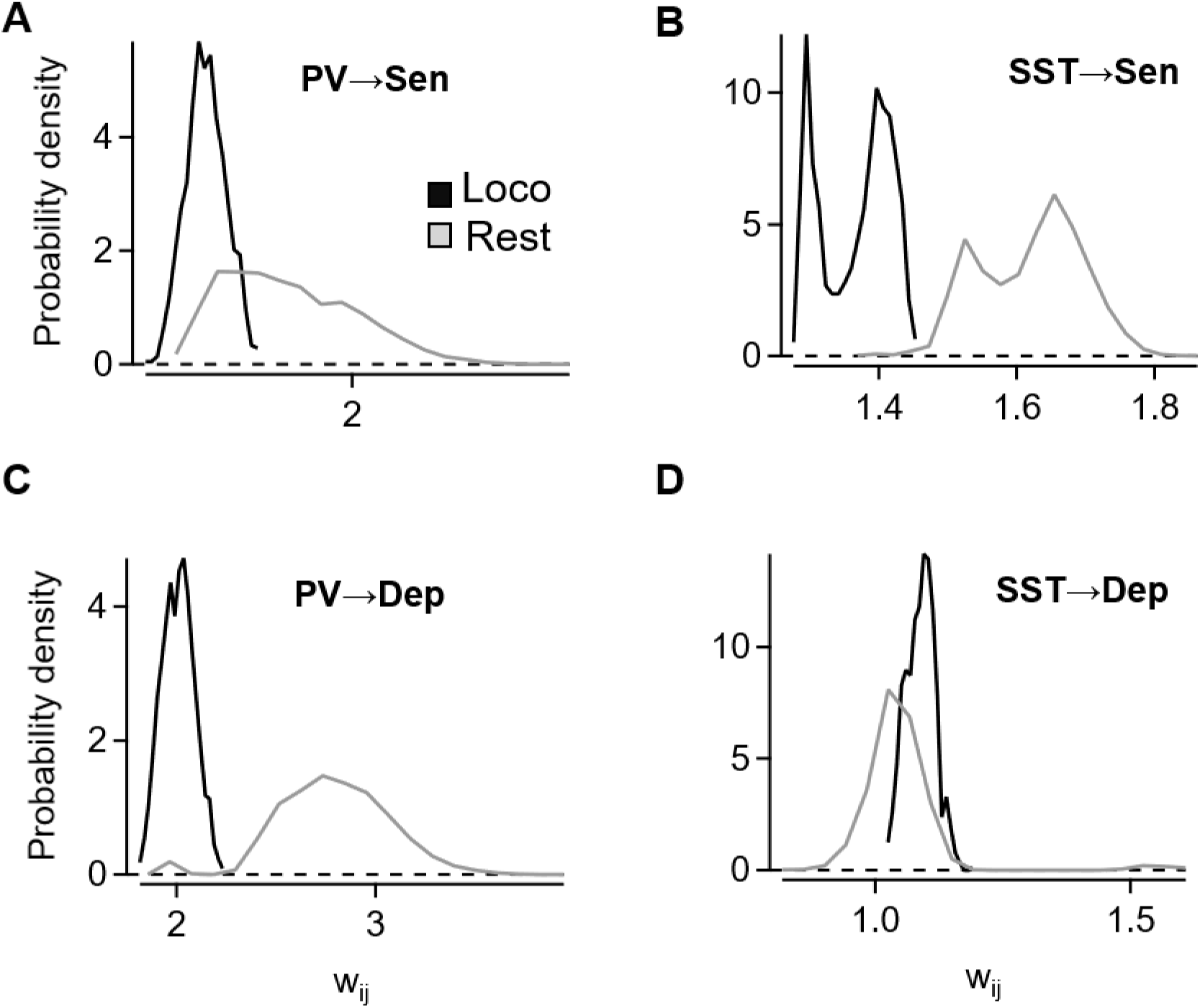
Parameter distributions across solutions for sensitizing and depressing PCs. Probability-density distributions of connection weights *w_iii_* across accepted model solutions at rest (grey) and during locomotion (black). **A-D**. Distributions are for PV→sensitizing PC (A), SST→sensitizing PC (B), PV→depressing PC (C), SST→depressing PC (E). The distributions shifted toward lower weights during locomotion, with the most prominent reductions occurring in PV input to depressing PCs and SST input to sensitizing PCs.

**Supplementary Figure 6.**
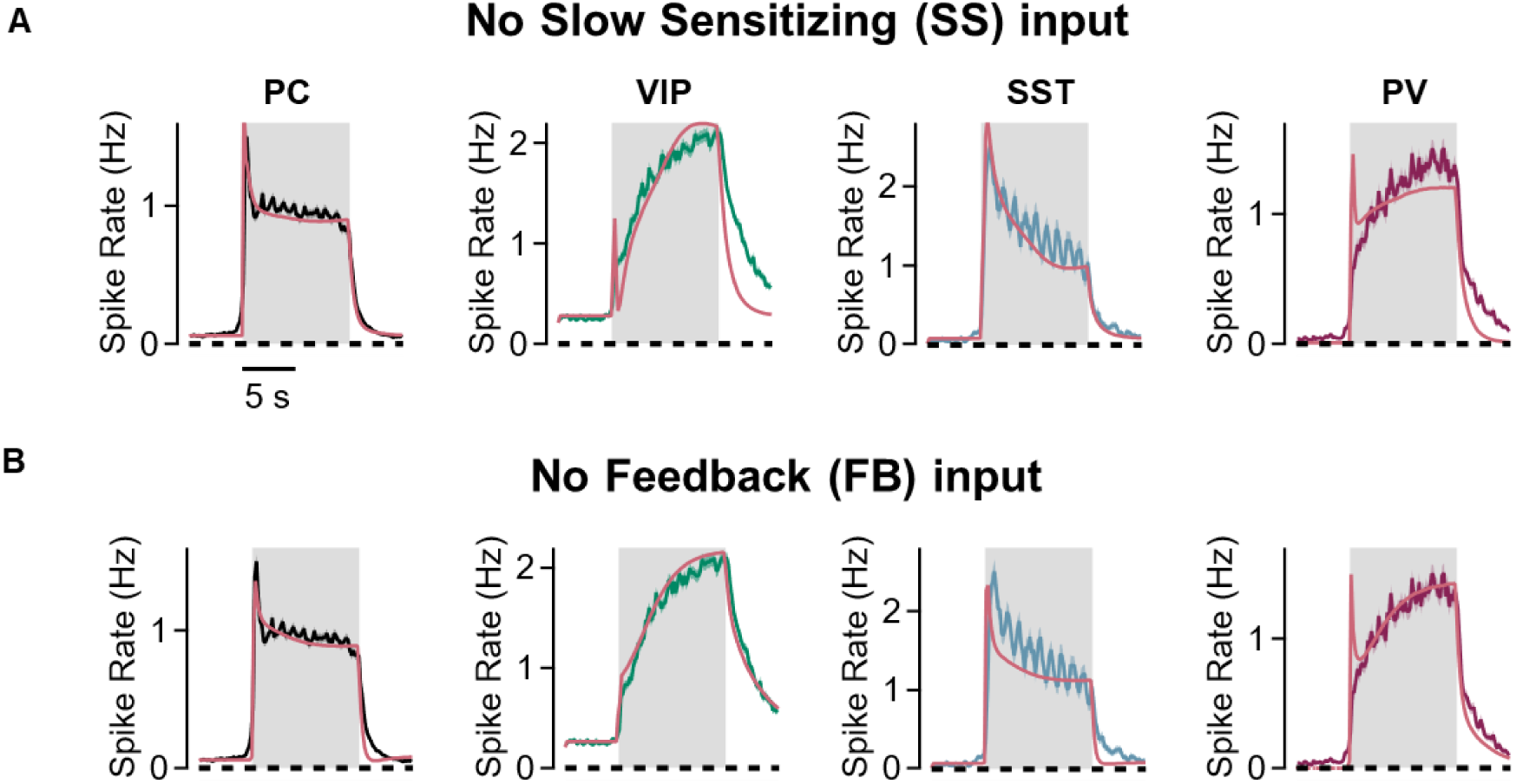
Removing external inputs impairs model performance. **A.** Average firing rates of PCs (black), VIPs (green), SSTs (blue) and PVs (dark red) during stimulus presentation during locomotion and their corresponding model fits after removal of the slow-sensitizing (SS) input. The solution shown is the one with the lowest RMSE among all solutions obtained with this reduced model (RMSE = 0.14), which was higher than the RMSE of the complete model (0.11). Note that VIP and PV dynamics are not properly captured. **B.** Same as in A after removal of the feedback (FB) input. The solution shown is the one with the lowest RMSE among all solutions obtained after FB removal (RMSE = 0.14), which was higher than the RMSE of the complete model (0.11). Note that SST and PV dynamics are not properly captured.

**Supplementary Table 1.** Locomotion-associated changes in the number of neurons responding to the visual stimulus.

| **Cell** | **Total**<br>**(N, per FOV)** | **Rest**<br>**responsive**<br>**(N, per FOV)** | **Locomotion**<br>**responsive**<br>**(N, per FOV)** | **Rest responsive**<br>**only**<br>**(N, per FOV)** | **Relative**<br>**increase during**<br>**locomotion** |
| --- | --- | --- | --- | --- | --- |
| **PC** | 78.6 ± 9.4 | 4.4 ± 1.1 | 27.8 ± 2.0*** | 0.5 ± 0.2 | 6.3 ± 1.7 |
| **SST** | 13.7 ± 1.6 | 5.1 ± 1.0 | 6.2 ± 0.7 | 0.5 ± 0.2 | 1.2 ± 0.3 |
| **PV** | 32.9 ± 5.4 | 10.1 ± 2.2 | 18.4 ± 1.6*** | 0.3 ± 0.2 | 1.8 ± 0.4 |
| **VIP** | 53.8 ± 7.2 | 11.7 ± 6.2 | 34.9 ± 3.2* | 1.3 ± 0.7 | 3.0 ± 1.6 |

From left: Total number of cells of each type imaged in a field of view (FOV) of 0.2 mm<sup>2</sup>. The number of cells generating a statistically significant response to the stimulus at rest (n = 11, 8, 16, 6 FOVs for PCs, SSTs, PVs and VIPs, respectively) and during locomotion (n = 55, 23, 35, 22). There was a significant increase in the number of responsive PCs and PVs (LMM: p < 0.001 in both) and VIPs (p < 0.05). Locomotion had no significant effect on the number of responsive SSTs. Many fewer cells were responsive at rest but not locomotion, indicating that most cells responsive at rest were also responsive during locomotion. Last column shows the relative change in the number of responsive neurons during locomotion compared to rest.

**Supplementary Table 2.** Dimensionality reduction criteria.

| Weight<sup>A</sup> | U<sub>Norm</sub><br>(1<sup>st</sup>)<sup>B</sup> | U<sub>Norm</sub><br>(2<sup>nd</sup>)<sup>C</sup> | % PC1<br>Contribution<sup>D</sup> | Model Status<sup>E</sup> |
| --- | --- | --- | --- | --- |
| W<sub>PC,PC</sub> / w_0 | 0.6 | 0.5 | 0.3 | Fixed |
| W<sub>PC,FF</sub> / w_1 | 0.6 | 0.5 | 0.6 | Fixed |
| W<sub>PC,SS</sub> / w_2 | 0.8 | 1.0 | 1.8 | Variable |
| W<sub>PC,PV</sub> / w_3 | 0.3 | 0.1 | 0.5 | Variable |
| W<sub>PC,SST</sub> / w_4 | 0.4 | 0.5 | 0.2 | Fixed |
| W<sub>PV,PC</sub> / w_5 | 0.2 | 0.3 | 2.0 | Variable |
| W<sub>PV,FF</sub> / w_6 | 0.8 | 0.7 | 2.4 | Variable |
| W<sub>PV,SS</sub> / w_7 | 0.4 | 0.5 | 0.1 | Fixed |
| W<sub>PV,PV</sub> / w_8 | 0.5 | 0.5 | 0.0 | Fixed |
| W<sub>PV,SST</sub> / w_9 | 0.4 | 0.5 | 0.3 | Fixed |
| W<sub>SST,PC</sub> / w_10 | 0.1 | 0.0 | 3.3 | Variable |
| W<sub>SST,FB</sub> / w_11 | 0.4 | 0.5 | 0.1 | Fixed |
| W<sub>SST,VIP</sub> / w_12 | 0.7 | 0.3 | 0.2 | Fixed |
| W<sub>VIP,PC</sub> / w_13 | 0.0 | 0.0 | 4.0 | Variable |
| W<sub>VIP,SST</sub> / w_14 | 0.9 | 0.8 | 2.7 | Variable |
| W<sub>VIP,SS</sub> / w_15 | 1.0 | 1.0 | 4.0 | Variable |
| W<sub>PC,FB</sub> / w_16 | 0.5 | 0.5 | 0.1 | Fixed |
| W<sub>PV,FB</sub> / w_17 | 0.4 | 0.5 | 0.6 | Fixed |
| W<sub>VIP,FB</sub> / w_18 | 0.9 | 0.8 | 3.3 | Variable |

**A.** Name of the model parameter as presented in the manuscript and in the code (text/code). **B-C.** Effect sizes between accepted locomotion and rest solutions, quantified using the normalized Mann–Whitney U statistic (Vargha and Delaney 2000), following the first reduction round, in which nine weights were fixed (B), and the second round, in which ten weights were fixed (C). Blue indicates an increase with locomotion, red a decrease, and white no meaningful change. **D.** Contribution of each parameter to the variance captured by the first principal component of the differences between locomotion and rest weights in the original unrestricted solutions. **E.** Final decision to fix each parameter across conditions or retain it as variable.

**Supplementary Table 3.** Average response amplitudes in resting and active states.

|  | **Rest**<br>(Amp<br>in Hz) | **Locomotion**<br>(Amp in Hz) | **Relative**<br>**Change<sup>c</sup>** |
| --- | --- | --- | --- |
| **PC** | 0.57 | 0.96 | 0.70 |
| **SST** | 1.52 | 1.45 | -0.05 |
| **PV** | 0.93 | 1.16 | 0.25 |
| **VIP** | 0.67 | 1.63 | 1.43 |

Note that activity in external inputs was not directly measured. We therefore kept these constant between rest and locomotion, so that when fitting the model changes in state were reflected only in changes in connection weights within layer 2/3.

**Supplementary Table 4.**
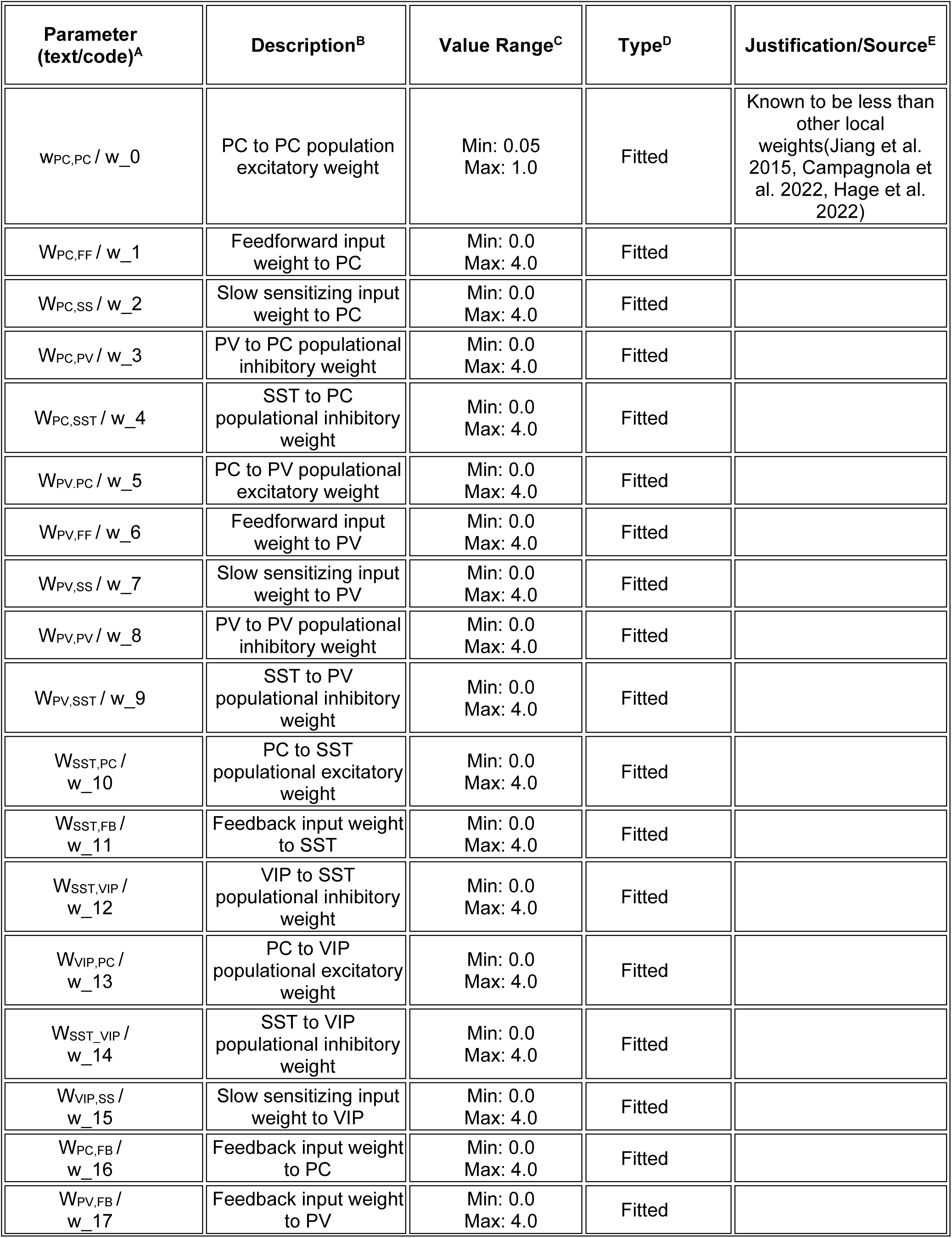
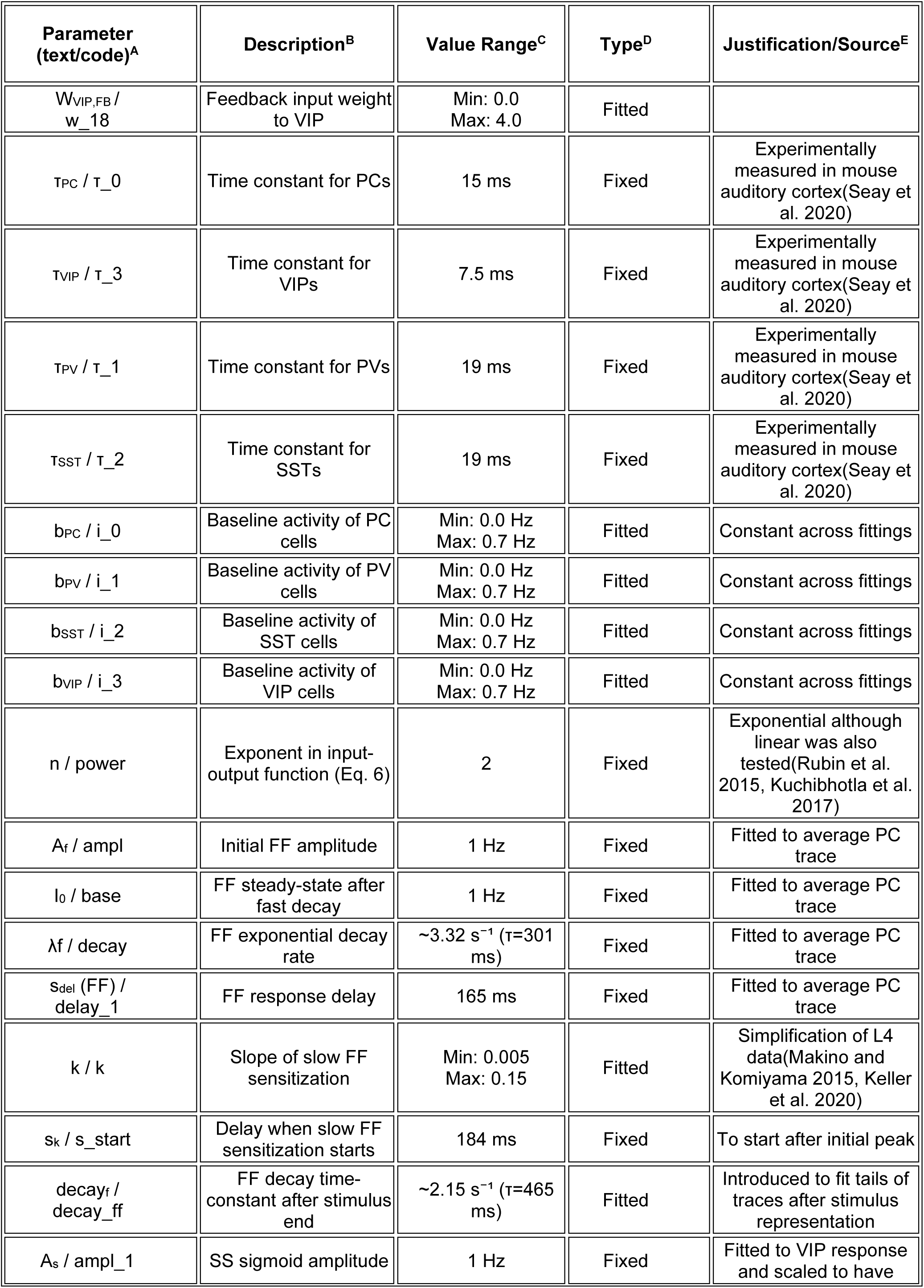
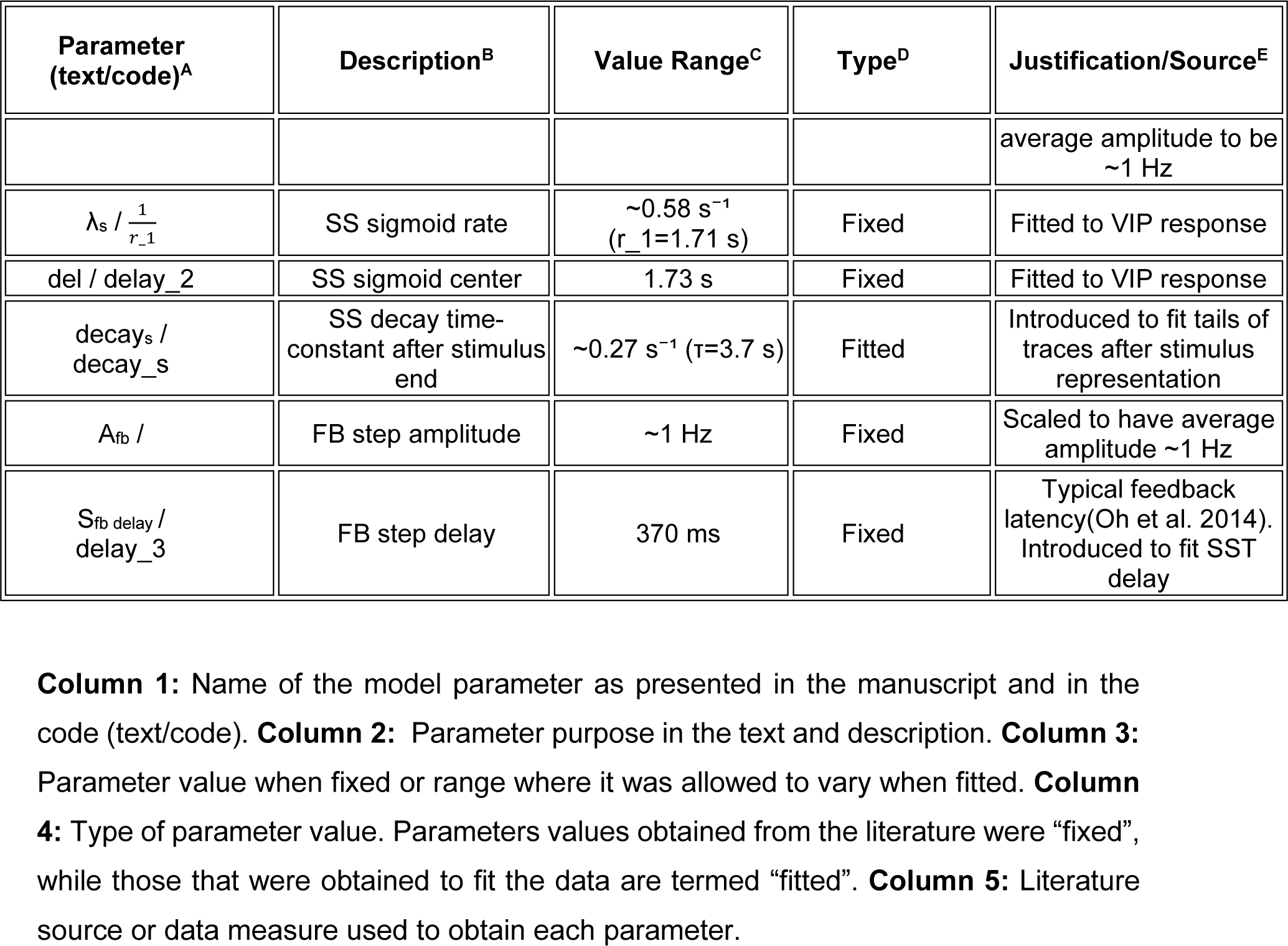
List of parameters used to fit the model.

